# Jan and mini-Jan, a model system for potato functional genomics

**DOI:** 10.1101/2024.12.10.627817

**Authors:** Haoyang Xin, Luke W. Strickland, John P. Hamilton, Jacob K. Trusky, Chao Fang, Nathaniel M. Butler, David S. Douches, C. Robin Buell, Jiming Jiang

**Affiliations:** Department of Plant Biology, Michigan State University, East Lansing, Michigan 48824, USA; Center for Applied Genetic Technologies, University of Georgia, Athens, Georgia 30602, USA; Department of Crop and Soil Sciences, University of Georgia, Athens, Georgia 30602, USA; Department of Horticulture, University of Wisconsin-Madison, Madison, Wisconsin 53706, USA; United States Department of Agriculture-Agricultural Research Service, Vegetable Crops Research Unit, Madison, Wisconsin 53706, USA; Department of Plant, Soil, and Microbial Sciences, Michigan State University, East Lansing, Michigan 48824, USA; Michigan State University AgBioResearch, East Lansing, Michigan 48824, USA; Institute of Plant Breeding, Genetics and Genomics, University of Georgia, Athens, Georgia 30602, USA; The Plant Center, University of Georgia, Athens, Georgia 30602, USA; Department of Horticulture, Michigan State University, East Lansing, Michigan 48824, USA

**Author notes:** These authors contributed equally to this work (Haoyang Xin, Luke W. Strickland). Yazhouwan National Laboratory, Sanya, Hainan Province, China, 572024 (Chao Fang). We dedicate this paper to Dr. Shelley Jansky, who has devoted her entire career to potato genetics and breeding. Dr. Jansky is credited for developing several key germplasm stocks, including M6 and DMF5163, the progenitor lines for Jan and mini-Jan. ‘Jan’ is named from ‘Jansky’.

## Abstract

Potato (*Solanum tuberosum*) is the third most important food crop in the world. Although the potato genome has been fully sequenced, functional genomics research of potato lags relative to other major food crops due primarily to the lack of a model experimental potato line. Here, we present a diploid potato line, ‘Jan’, which possesses all essential characteristics for facile functional genomics studies. Jan has a high level of homozygosity after seven generations of self-pollination. Jan is vigorous and highly fertile with outstanding tuber traits, high regeneration rates, and excellent transformation efficiencies. We generated a chromosome-scale genome assembly for Jan, annotated genes, and identified syntelogs relative to the potato reference genome assembly DMv6.1 to facilitate functional genomics. To miniaturize plant architecture, we developed two “mini-Jan” lines with compact and dwarf plant stature using CRISPR/Cas9-mediated mutagenesis targeting the *Dwarf* and *Erecta* genes related to growth. Mini-Jan mutants are fully fertile and will permit higher-throughput studies in limited growth chamber and greenhouse space. Thus, Jan and mini-Jan provide an outstanding model system that can be leveraged for gene editing and functional genomics research in potato.

## Introduction

Cultivated potato (*Solanum tuberosum*, 2n = 4x = 48) is the third-most important global food crop for human consumption (Devaux et al., 2020), with approximately 375 million tons produced from nearly 18 million hectares in 2022 alone (http://www.fao.org/). French fries and potato chips are among the most popular snack foods in the world, especially in developed countries. Unlike most major crops, development of new potato cultivars has been hindered by characteristics inherent to its biology, especially its highly heterozygous outcrossing autotetraploid genome, clonal propagation nature, and sensitivity to inbreeding depression due to high genetic load. Modern potato cultivars developed in the United States after the 1970s showed similar yield potential as those developed in the 19^th^ century (Douches et al., 1996), despite high yield being a top breeding goal for most potato breeding programs. Recent genome sequencing and transcriptomic analysis of several tetraploid potato cultivars have revealed extensive allelic diversity and numerous dysfunctional or deleterious alleles (Hoopes et al., 2022; Mari et al., 2024; Sun et al., 2022). These genomic features of tetraploid potato have hindered breeders’ efforts to reduce genetic load underlying the lack of yield increase over 100 years of traditional breeding.

After a century-long struggle with tetraploid potato, the research community has initiated a diploid inbred-based system for potato breeding (Bethke et al., 2022; de Vries et al., 2023; Jansky et al., 2016). The value of diploid potato (2n = 2x = 24), including wild diploid species and haploids (or “dihaploids”) derived from tetraploid cultivars, has long been recognized. In fact, the first genetic linkage maps of potato were generated with diploid populations (Bonierbale et al., 1988; Gebhardt et al., 1989). Identification and cloning of key genes in potato has mostly relied on genetics and genomics research of diploid potato (Ballvora et al., 2002; Eggers et al., 2021; Kloosterman et al., 2013; Ma et al., 2021; Song et al., 2003). However, most currently available diploid potato species or breeding lines share genetic and genomic characteristics that are not desirable for functional genomics studies, including high heterozygosity and self-incompatibility. Self-compatible and homozygous accessions of diploid species have been reported, including *Solanum verrucosum* (Hosaka et al., 2022) and *Solanum chacoense* (Jansky et al., 2014). However, these genotypes are often highly recalcitrant to regeneration and are rarely used for transgenic research (Duangpan et al., 2013). Although an increasing number of diploid breeding lines have been evaluated in recent years (Achakkagari et al., 2022; Alsahlany et al., 2021; Hosaka and Sanetomo, 2020; Jayakody et al., 2022; Jayakody et al., 2023), the potato research community is still in need of a model line for functional genomics studies. Such a line should be vigorous, self-compatible, with excellent tuber traits, and most importantly, can be readily transformed using Agrobacterium.

Here, we describe the development of ‘Jan’, a diploid potato line derived from a hybrid between *S. tuberosum* Group Phureja clone DM1-3 516 R44 (hereafter referred to as DM) (Pham et al., 2020; The Potato Genome Sequencing Consortium, 2011) and M6, a self-compatible accession of *S. chacoense* (Jansky et al., 2014; Leisner et al., 2018). Jan was self-pollinated for seven generations and thus, reached a high level of homozygosity. Jan is highly fertile with outstanding tuber traits. More importantly, Jan can be readily regenerated and transformed Agrobacterium. Thus, Jan combines the most desirable traits from both parents, including the vigor and fertility from M6 and the tissue culture and transformation amenability from DM. We generated a chromosome-scale genome assembly of Jan and annotated it for protein coding genes to facilitate gene identification and mutational research. We also developed several “mini-Jan” mutants with compact and dwarf plant statue using CRISPR/Cas9-mediated mutagenesis of two genes related to growth. The mini-Jan lines are fully fertile and will permit increased capacity in growth chamber and greenhouse studies. Thus, Jan and mini-Jan provide a model system for functional genomics and molecular genetics research in potato.

## Results

### Morphology and fertility of Jan

We identified the self-compatible clone DMF5163 as a starting material to develop a model diploid potato line. DMF5163 was derived from a cross between DM and M6 (Endelman and Jansky, 2016) and was selfed for five generations in the greenhouse. DMF5163 was self-pollinated for two additional generations under growth chamber conditions. A single F7 individual (DMFJ7), named ‘Jan’, was selected from the growth chamber population largely based on its vigor and fertility. Jan exhibits vigorous growth and produces abundant flowers under standard growth chamber and greenhouse conditions (**Figure 1**). Jan has a compact plant structure that reaches an average height of 57 cm in the growth chamber (**Figure 1a**) and is fully mature after 105 days in the greenhouse. Jan tubers are round with a cream skin color and shallow eyes and are relatively uniform in size (**Figure 1e**). Jan produced an average of 33 tubers per plant with a yield of 378 g in the greenhouse. Jan is self-compatible and produces abundant viable pollen (**Figure S1a**). Approximately 90% of the pollen showed normal I_2_-KI staining (Pedersen et al., 2004) (**Figure S1b**). Jan plants had a 94% fruit-setting rate upon self-pollination with berries generating an average of 65 seeds per fruit (**Figure S1c**).

**Figure 1.**
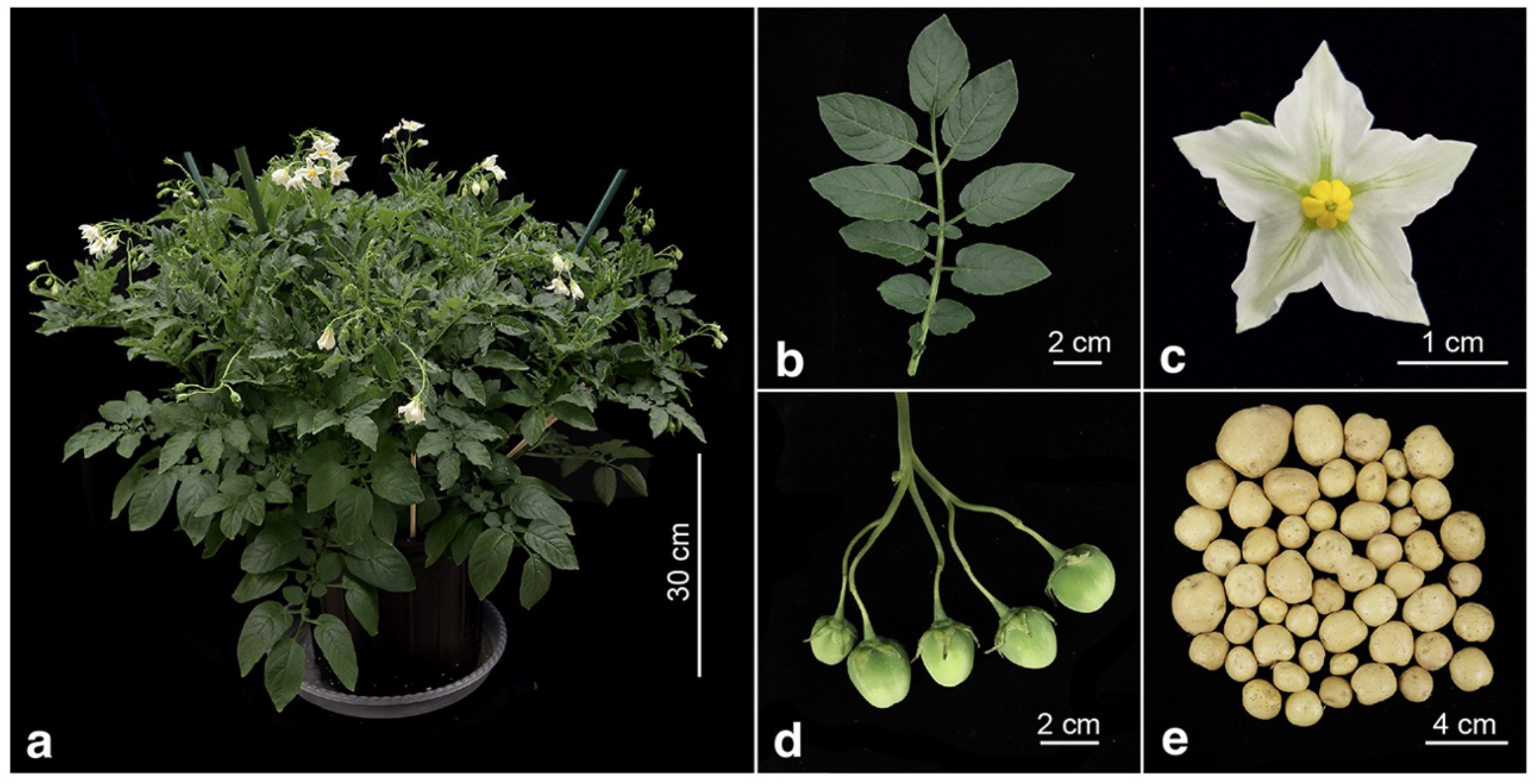
Phenotypic characteristics of Jan. (**a**) Plant architecture. (**b**) Leaflets from a single compound leaf. (**c**) Flower. (**d**) Fruits. (**e**) Tubers from a single plant grown in a growth chamber.

### Development of a reference genome of Jan

We developed a chromosome-scale genome assembly for Jan, an essential resource for Jan being used as a model functional genomics tool. We generated 1,927,473 sequence reads of 10 kb or longer using Oxford Nanopore Technologies (ONT) long read sequencing, totaling 53.1 Gb of sequence and representing ∼63X genome coverage (**Table S1**) that were assembled using Flye (Kolmogorov et al., 2019). Error correction of the draft assembly was performed using both ONT long reads and Illumina whole-genome shotgun reads (**Table S1**). Duplicative and short contigs were filtered out yielding an assembly of 717.2 total Mb from 953 contigs with an N50 length of 6.7 Mb. Using DM v6.1 (Pham et al., 2020) as the reference, the contigs were placed onto the 12 chromosomes resulting in final assembly of 717.2 Mb, of which, 708.0 Mb was scaffolded to the 12 potato chromosomes (**Table S2**). LTR assembly index (LAI) assessment of the assembly revealed a score of 13.28, indicative of a reference quality assembly (10 σ; LAI σ; 20) (Ou et al., 2018). Benchmarking Universal Single-Copy Orthologs (BUSCO) analysis (Manni et al., 2021) revealed 98.2% complete BUSCOs for the assembly (**Table S3**), indicative of a high-quality assembly. Analyses of whole-genome shotgun reads indicate the presence of some residual heterozygosity within the Jan genome (**Figure S2**) (Mapleson et al., 2017; Ranallo-Benavidez et al., 2020).

To annotate the Jan genome, we performed *de novo* repetitive sequence identification, revealing 65.8% repeat content (**Table S4**), similar to the repetitive sequence content determined previously for M6 (60.7%) (Leisner et al., 2018) and DM (66.8%) (Pham et al., 2020). Using the repeat-masked genome, protein-coding protein sequences were annotated using BRAKER with further refinement of the models using PASA (v2.5.2; (Campbell et al., 2006)) with RNA-sequencing and full-length cDNA sequences (**Table S1**). A total of 71,186 working gene models were annotated. Of these, 64,288 high confidence gene models were annotated from 35,985 loci; BUSCO analysis revealed 89.5% complete BUSCOs for the annotation (**Table S3)**.

We observed a high rate of syntelog mapping between the high-confidence gene models identified in Jan and its two parents (**Figure S3**), as well as those of other diploid potatoes, including DM1S1 (Jayakody et al., 2023) and *Solanum candolleanum* (http://spuddb.uga.edu/S_candolleanum_v1_0_download.shtml), and the non-potato *Solanum* species tomato (*Solanum lycopersicum*) (Hosmani et al., 2019) (**Figure S3**). In total, 25,943 syntelogs were identified between Janv1.1 and DMv6.1. As DM has served as the reference genome for potato since 2011 and the two parents of Jan differ in numerous traits, the high rate level of synteny identified in genes among Jan, DM, and M6 (**Dataset 1**) will facilitate effective gene identification and functional genomics assays based on Jan.

### Representation of the parental genomes in Jan

We used the chromosome-scale genome assemblies of DM and M6 to determine which parental alleles are represented in Jan. Sequence alignment of Jan against the genomes of DMv6.1 (Pham et al., 2020) and M6v5.0 (https://spuddb.uga.edu/M6_v5_0_download.shtml) in pairwise fashion revealed large sections of collinearity (**Figure S4**). One notable observation is the inheritance of the entire centromeric and pericentromeric regions from DM on chromosomes 1, 4, 9 (aside from an apparent inversion near the centromere), 10, and 11, and from M6 on chromosomes 3 and 7 (**Figure S4**). The inheritance of these regions of the remaining chromosomes is less clear.

To identify blocks of genomic sequence inherited from each parent in Jan, k-mers from the genome assembly of each parent were anchored to the Jan assembly and k-mer conservation between Jan and each parent was calculated in 100 kb windows (Aylward et al., 2023) (**Figure S5**). Windows with k-mer conservation differences greater than 15% were assigned to the parent with the higher conservation; windows with 15% or less k-mer conservation difference (i.e., high sequence conservation between DM and M6), were assigned as ambiguous inheritance. In total, of the 708 Mb scaffolded to the twelve chromosomes, Jan inherited 44.4% of its genomic sequence from DM and 31.6% from M6; 24.0% were ambiguous due to high sequence homology between the two parental genomes (**Figure 2**).

**Figure 2.**
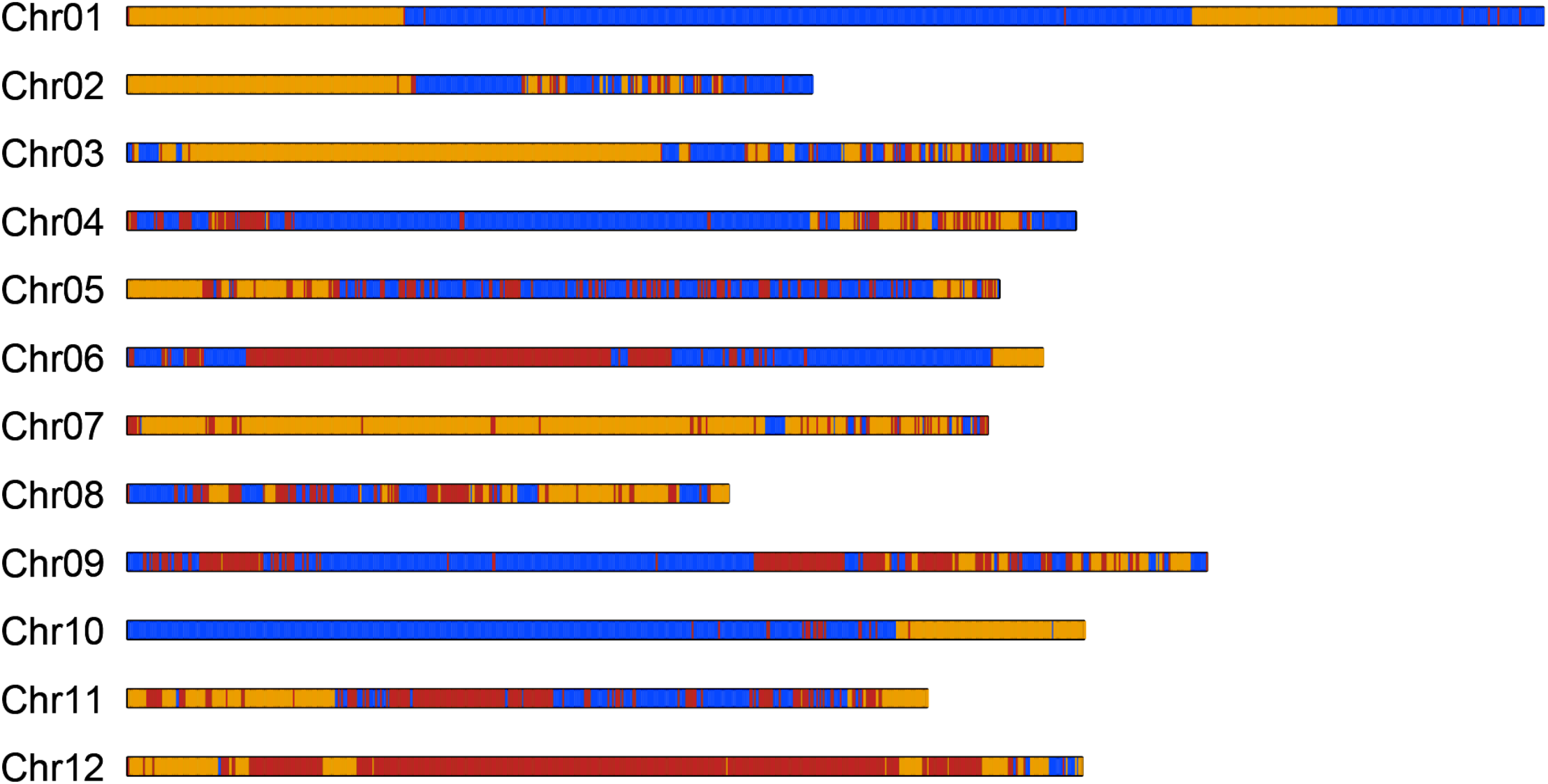
Allelic representation of DM and M6 in the Jan genome. Blocks of genomic sequence are in 100 kb resolution and color-coded by its parental origin: DM (blue), M6 (gold), or ambiguous (red) due to high sequence conservation between DM and M6.

### Putative parental genes relevant to the distinct phenotypes of Jan

The annotated genes of Jan were classified as DM or M6 alleles based on the k-mer conservation classification. Of the 35,700 genes placed onto the twelve chromosomes, 15,056 (42.2%) and 13,436 (37.6%) genes were inherited from DM and M6, respectively. The remaining 7,208 (20.2%) were ambiguous. Interestingly, genes inherited from each parent were differentially enriched in specific biological processes and molecular functions in Gene Ontology (GO) term analysis. Genes inherited from DM were enriched for “catabolic process”, “manganese ion binding”, and “structural constituent of chromatin”; while those from M6 were enriched in “response to hormone”, “organic acid transport”, “recognition of pollen”, and “anatomical structure development” (**Table S5**).

Jan is vigorous and highly fertile, resembling M6. In contrast, DM is a very weak plant and male sterile. To explore the genetic basis of these distinct parental traits, we investigated the significant enrichment of inherited genes and their expression in six developmentally important tissues: young (immature) leaves, flower buds, open (mature) flowers, stolons, small tubers, and roots. Of the 47 genes inherited from M6 and annotated under “recognition of pollen” (GO:0048544, *p*-value 1.3e-08) category, there are 42 receptor-like kinases (RLKs), including 20 lectin RLKs (Lec-RLKs). Previous studies have showed the important roles of Lec-RLKs in determining male fertility by regulating pollen exine assembly and pollen aperture development in both *Arabidopsis thaliana* and rice (Peng et al., 2020; Wan et al., 2008). A total of 18 of these RLKs (six of which are Lec-RLKs) exhibit moderate expression (5-40 transcripts per million (TPM) in floral tissues, suggesting a potential contribution in Jan’s fertility. Furthermore, four phospholipase A2 family genes annotated under “organic acid transport” (GO:0015849, *p*-value 6.0e-04) are included in the 27 significant genes inherited from M6, and three of them are moderately to highly expressed in floral tissues (13-85 TPM) (**Dataset 2**). It has been demonstrated in *A. thaliana* that phospholipases are essential for proper pollen development, since RNA interference (RNAi) knockdowns of these genes result in pollen lethality (Kim et al., 2011).

Both M6 and Jan are highly vigorous. Interestingly, we identified several M6-derived genes related to plant growth hormone signaling and response. The 46 significant genes inherited from M6 and annotated under the “response to hormone” (GO:0009725, *p*-value 3.5e-05) category include 38 auxin response factor and/or SAUR-like auxin-responsive protein family genes, 18 of which are found in tandem or are organized as gene clusters on four different chromosomes. Of these, 25 of the 38 genes are moderately to highly expressed (5-209 TPM) in at least one analyzed tissue, and more often in multiple tissues. Also included in this group are five major latex protein (MLP)-like genes. MLP-like genes are known to promote vegetative growth through response to *cis*-cinnamic acid (Guo et al., 2011). One of these genes (Soltu.Jan_v1.1.09G030670.1) exhibits a very high expression (80-1313 TPM) in all tissues analyzed. Particularly intriguing are the 94 significant genes inherited from M6 and annotated under the “anatomical structure development” (GO:0048856, *p*-value 5.9e-03) category. This includes eight plant-specific YABBY transcription factor (TF) family genes, six of which are moderately to highly expressed (6-185 TPM) and display preferential expression in above ground tissues (young leaf, flower bud, open flower). YABBY TFs are known to play an important in determination of abaxial cell fates in leaves and flowers (Siegfried et al., 1999).

Like DM and unlike M6, Jan can be regenerated in tissue culture efficiently and is readily transformable. Of the 266 significant genes inherited from DM and annotated under the “catabolic process” (GO:0009056, *p*-value 8.3e-06) category, two are arginine decarboxylase (ADC) genes. Arginine decarboxylation by ADC enzymes initiates the biosynthesis of the polyamine putrescine (Martin-Tanguy, 2001), and plays an important role in callus induction. Previous studies showed a positive correlation between ADC activity (presumably leading to increased putrescine content) and callus growth (Burtin et al., 1989; Hao et al., 2005). Furthermore, an increase in putrescine was shown to contribute to higher shoot regeneration frequencies in tissue culture (Bais et al., 2000; Fazilati and Forghani, 2015). Both identified ADC genes exhibit moderate to very high expression (29-538 TPM) in all tissues analyzed in Jan, with some of the higher expression values detected in roots of tissue culture plantlets (164, 176 TPM). Therefore, the high expression of DM-inherited ADC genes could lead to an increased supply of putrescine and successful regeneration observed in Jan.

We identified a globally expressed gene (6-11 TPM) encoding a SWIB complex BAF60b domain-containing protein (Soltu.Jan_v1.1.01G050000.1), one of the 50 significant genes inherited from DM and annotated under “manganese ion binding” (GO:0030145, *p*-value 8.9e-12) category. RNAi knockdown of the *A. thaliana* SWIB complex ortholog CHC1 resulted in reduced callus formation in tissue culture and consequently reduced *Agrobacterium*-mediated transformation rates (Crane and Gelvin, 2007). In addition, of the 29 significant genes inherited from DM and annotated under “structural constituent of chromatin” (GO:0030527, *p*-value 5.3e-04) category are 22 moderately to very highly expressed (5-2390 TPM) histone superfamily protein genes, including three histone H2A genes. Previous studies revealed a crucial role of H2A and other histone-associated proteins in the establishment of *Agrobacterium*-mediated transformation efficiency in *A. thaliana* (Yi et al., 2006; Yi et al., 2002; Zhu et al., 2003). Insertion of a T-DNA in the 3’ UTR of an H2A gene resulted in severely reduced transformation rates, while overexpressing of this H2A gene doubled the transformation rates (Mysore et al., 2000). It should be noted that Jan also inherits six histone H2A genes from M6; however, the H2A genes inherited from DM displayed overall higher expression (115-357 TPM) in the roots of tissue culture plantlets compared to the H2A genes inherited from M6 (10-235 TPM). These results support more substantial contributions from the DM-inherited genes to Jan’s tissue culture regeneration phenotypes.

### Regeneration and transformation of Jan

An efficient regeneration and transformation system is essential for functional plant genomics research. To establish a robust genetic transformation system, we first evaluated the regeneration efficiency of Jan using a similar method previously developed for DM (Paz and Veilleux, 1999). We cultured internode explants derived from 4-week-old tissue culture plantlets on pre-culture medium for three days. Internodes were then transferred to regeneration medium for approximately one month. We determined the regeneration rate, defined as the percentage of explants that developed shoots. Jan showed an average regeneration efficiency of 89.7% from three independent experiments (**Table 1**), supporting Jan’s inheritance of its regeneration efficiency from DM.

**Table 1.** Shoot regeneration efficiency of Jan.

| Replicates | No. of explants | No. of regeneration | Regeneration efficiency (%)<sup>a</sup> |
| --- | --- | --- | --- |
| 1 | 32 | 29 | 90.6 |
| 2 | 30 | 27 | 90.0 |
| 3 | 35 | 31 | 88.5 |

<sup>a</sup>Number of regenerated plants divided by the total number of explants × 100%

CRISPR/Cas9-based gene editing experiments were used to determine the transformation and gene editing efficiency of Jan. To this end, we targeted a uridine diphosphate glucosyltransferase gene (Soltu.Jan-1.1.05G025420.1) for mutagenesis. We designed a single guide RNA (gRNA_6) to target this gene. The gRNA was driven by the Arabidopsis U6 promoter, while the Cas9 gene was expressed under the Cestrum Yellow Leaf Curling Virus (CmYLCV) promoter (Stavolone et al., 2003) (**Figure 3a**). Starting with 151 explants, we achieved a transformation efficiency of 15.9% (the number of transgene positive events divided by the total number of explants), with 24 out of the 116 regenerated plants (20.7%) confirmed to be transgenic by PCR detection of the kanamycin resistance gene (**Table 2**). A total of 17 of the 24 transgenic plants (70.8%) showed editing through Sanger sequencing followed by Inference of CRISPR Edits (ICE) analysis, with 14 having editing scores greater than 10% (Conant et al., 2022).

**Figure 3.**
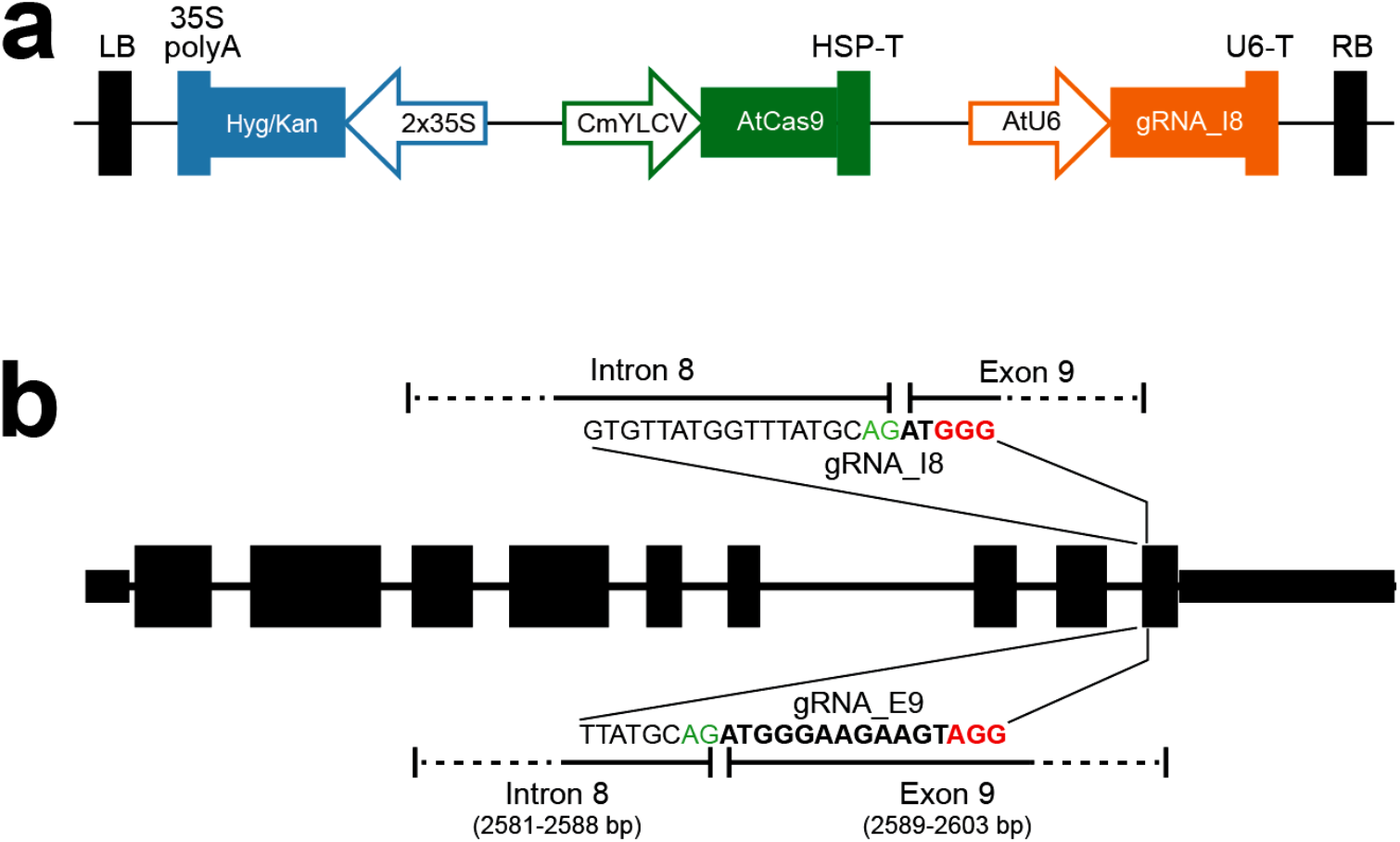
Diagrams of gRNAs and constructs for CRISPR/Cas9 experiments targeting the *StD* gene. (**a**) Illustration of the T-DNA region of the CRISPR/Cas9 construct. (**b**) Sequences and positions of the two gRNAs targeting the *StD* gene. Green color highlights “AG” represent the 3’ splicing site within intron 8. PAM sequences are highlighted in red. Bold letters represent sequence from exon 9. (**b**) Illustration of the T-DNA region of the CRISPR/Cas9 construct.

**Table 2.**
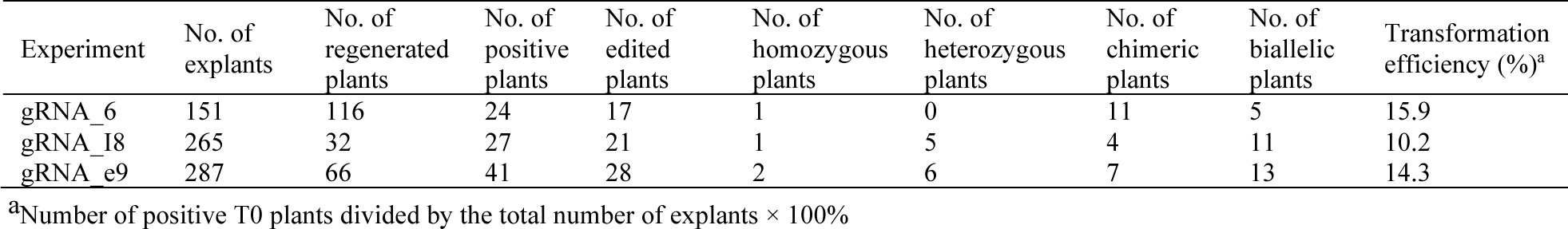
Transformation efficiency of Jan.

### Development of mini-Jan^D^

A miniaturized morphotype would permit a higher density of plants to be grown in a limited growth space and improve the capacity to perform functional genomics. Thus, we intended to develop a dwarf mutant of Jan, named “mini-Jan”. In tomato, mutation of the *Dwarf* or *D* gene resulted in the miniaturization of the Micro-Tom cultivar (Marti et al., 2006). In order to minituraize Jan, we targeted the *D* gene ortholog in Jan (Soltu.Jan_v1.1.02G031560.1) in Jan. In Micro-Tom, a single base mutation (A to T) of the 3’ <u>A</u>G splicing site of intron 8 in the *D* gene (**Figure 3b**) causes mis-splicing and truncation of the resulting protein (Marti et al., 2006). We attempted to recreate this mutation in Jan by using two different single 20-bp gRNAs spanning intron 8 and exon 9 of the *StD* gene. The cleavage site of gRNA_i8 was positioned at the 3’ AG splicing site and the resulting mutations of the AG site in Jan should mimic the mutation associated with Micro-Tom (**Figure 3b**). gRNA_e9 was targeted to exon 9 to generate proteins that would potentially be truncated within the last exon (**Figure 3b**). We generated 49 T0 plants with targeted mutations. We achieved a transformation efficiency of 10.2% in experiments using the gRNA_i8 construct; 27 of the 32 regenerated events (84.4%) were confirmed to be transgenic (**Table 2**). Among the transgenic plants, 52.4% displayed biallelic mutations. For the gRNA_e9 construct, 41 out of 66 regenerated plants (62.1%) were transgenic, resulting in an overall transformation efficiency of 14.3% (**Table 2**). Within this group, 46.4% were biallelic mutants. Combining the results from both experiments, the average transformation efficiency was 12.3% (**Table 2**), with an average biallelic or homozygous mutation rate of 55.4%.

We selected four representative edited lines (i8-2, 29-2, i8-1, e9-1) (**Figure 4a**) for in-depth analysis.

(1) i8-2: both *StD* alleles were mutated, with a 7 bp and 2 bp deletion in intron 8, respectively (**Figure 4c**). The 2 bp deletion did not affect the AG slicing site, and produces a normal transcript, which was detected in analysis of transcripts sequenced from i8-2 (**Figure 4c**). In contrast, the second allele with a 7 bp deletion lost the AG site, which resulted in two different types of short transcripts due to premature stop codons. The short transcripts encode two truncated proteins losing 24 and 25 amino acids (aa), respectively (**Figure 4c**), resembling the mutations described in Micro-Tom (Marti et al., 2006). The morphology of i8-2 plants are highly similar to Jan (**Figure 4a**), likely due to the presence of normal *StD* transcripts in this mutant.
(2) e9-2: both *StD* alleles were mutated, with a 1 bp deletion and 1 bp insertion within exon 9, respectively (**Figure 4d**). Both mutated alleles resulted in a shortened transcript caused by a premature stop codon. The two short cDNAs encode two truncated proteins that lose 21 aa and 25 aa, respectively (**Figure 4d**). The e9-2 plants show a reduced height, and a more condensed form compared to Jan. Leaves from e9-2 plants are slightly wrinkled and show a darker pigmentation compared to Jan (**Figure 4a**).
(3) i8-1: both *StD* alleles were mutated, with 3 bp deletion and 1 bp insertion within intron 8, respectively. These mutations resulted in the AG splicing site being lost or mutated in both alleles (**Figure 4e**). Two different resulting transcripts were detected from i8-1. Both transcripts can be derived from either one of the two mutated alleles. The two transcripts encode two truncated proteins that lose 24 aa and 25 aa, respectively (**Figure 4e**). i8-1 plants show a semi-dwarf phenotype with leaves exhibiting a darker tone than Jan, although not as pronounced as i9-2. The leaves also showed a subtle crinkled appearance. The stem of i8-1 was moderately thicker than Jan. The stem of i8-1 was moderately thicker than Jan, characteristic of a dwarfed phenotype. The i8-1 plants have a reduced height and a bushier growth habit (**Figure 4a**). Inflorescences were drastically shorter, lacking any noticeable elongation. Both flowers and fruits of i8-1 are reduced in size compared to Jan. These characteristics suggest that i8-1 most closely resembles the Micro-Tom phenotype among the Jan mutants analyzed. Tissue culture seedlings of i8-1 showed pronounced short internodes and dwarf phenotypes (**Figure 5a**). The i8-2 mutant was named as mini-Jan^D^.
(4) e9-1: both *StD* alleles lost 1 bp within exon 9, leading to the loss of 25 amino acids (**Figure 4f**). e9-1 plants display a strong brassinosteroid deficiency symptoms described in other plant species, including tomato (Bishop et al., 1999) (**Figure 4a**). e9-1 leaves were a notably darker green with a texture reminiscent of crumpled paper. Compared to Jan, e9-1 leaves were shorter in length and took on a more rounded shape. e9-1 plants were significantly dwarfed, with a robust stem diameter (**Figure 4a**). e9-1 plants were completely sterile, thus, are not useful for any further application.

**Figure 4.**
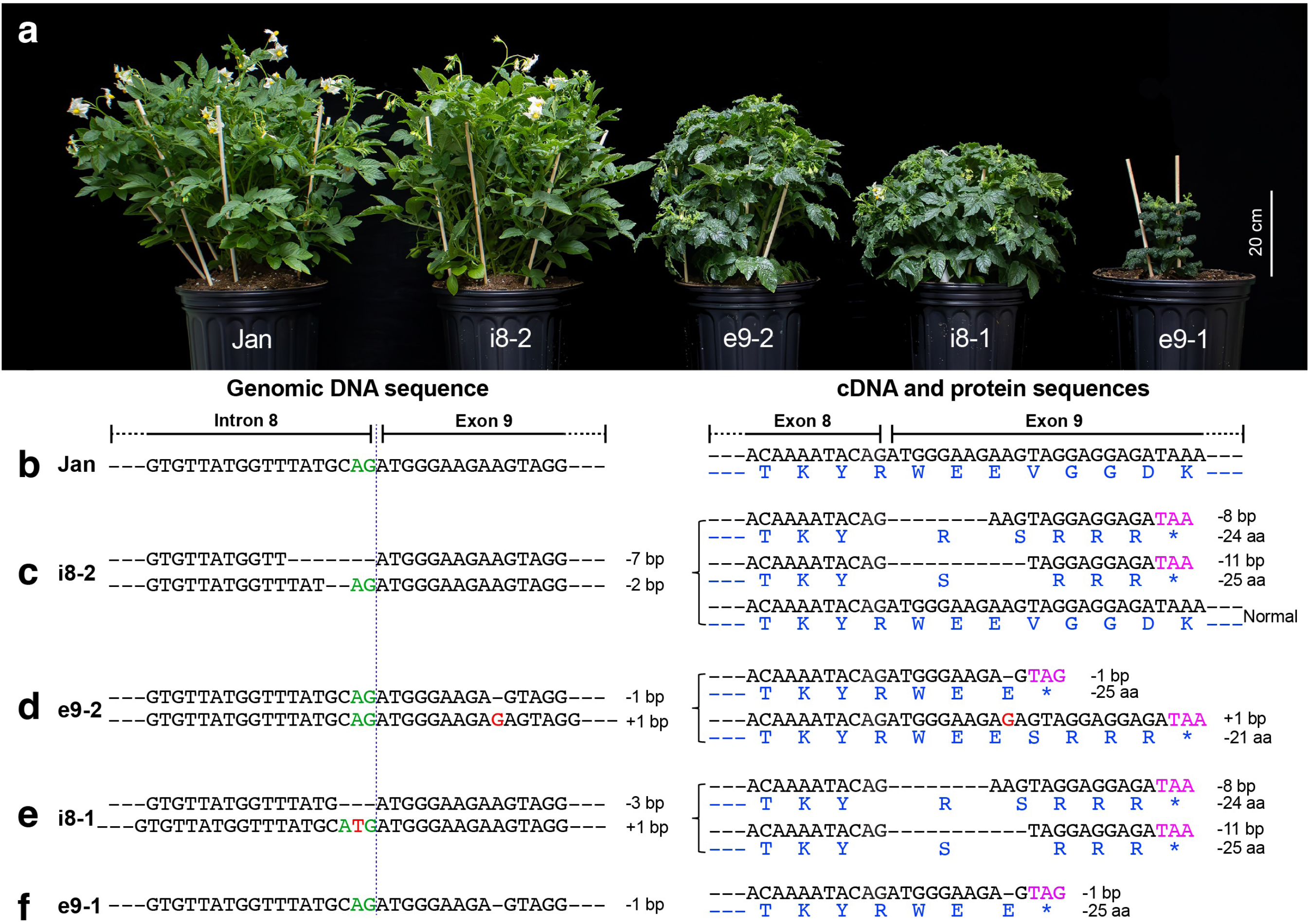
Genomic composition and phenotype of mini-Jan mutants from mutagenesis of the *StD* gene. (**a**) A single plant of Jan and four T0 mutants at 48 days after planting in a growth chamber. (**b-f**) Genomic DNA sequences, cDNA sequences, and predicted protein sequences of Jan (**b**), mutant i8-2 (**c**), mutant e9-2 (**d**), mutant i8-1 (**e**), and mutant e9-1 (**f**). The pre-mature stop codons are marked by magenta. The splicing AG sites are marked by green. The predicted protein sequences are in blue. The vertical blue line separates exon 9 from intron 8 sequence.

**Figure 5.**
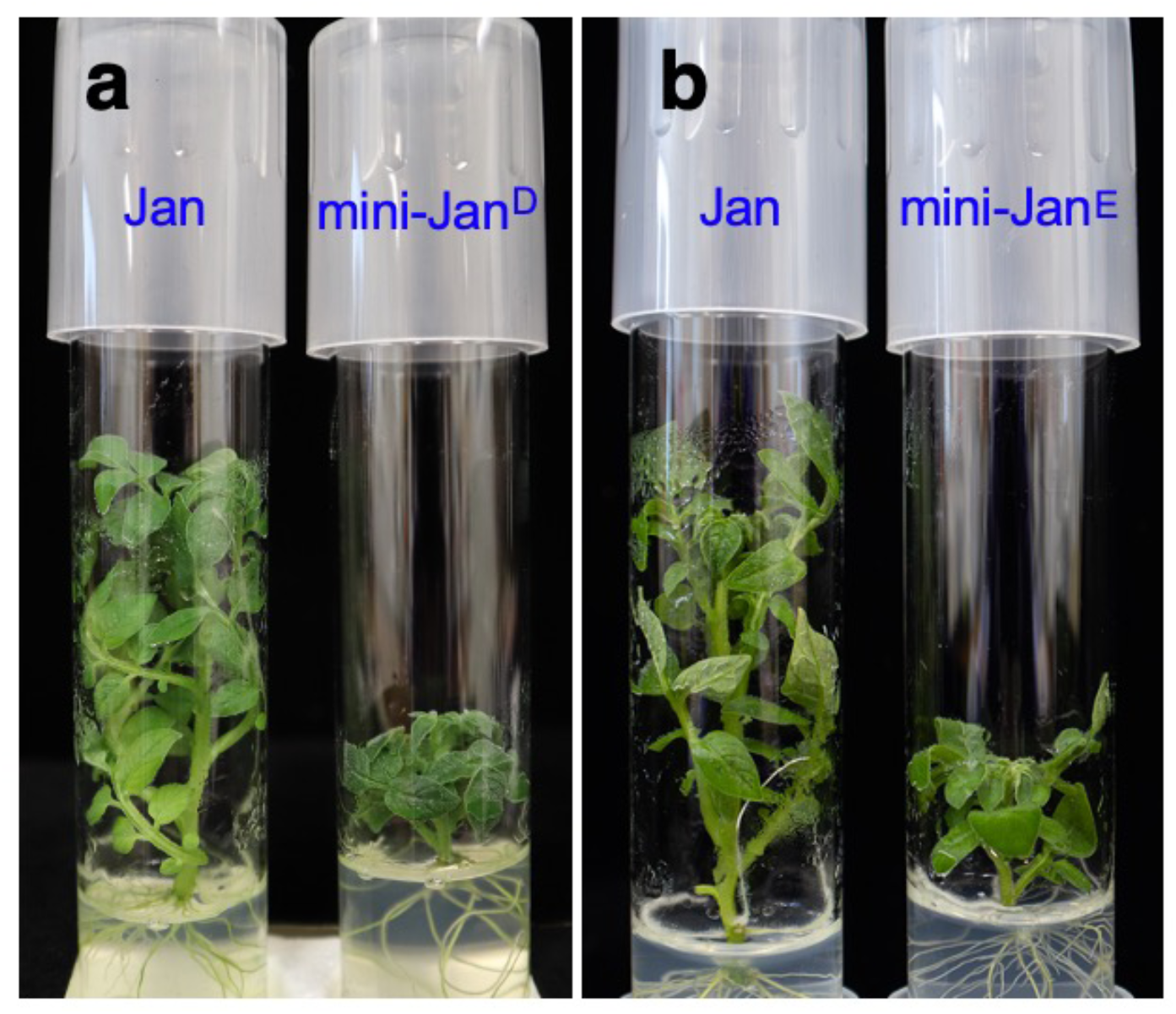
The phenotypes of tissue culture plants of Jan and mini-Jan. (**a**) Tissue culture plants of Jan and mini-Jan^D^ after 25 days of culture. (**b**) Tissue culture plants of Jan and mini-Jan^E^ after 20 days of culture. Note: both mini-Jan^D^ and mini-Jan^E^ show a pronounced dwarf phenotype compared to the wild type Jan.

### Development of mini-Jan^E^

We also explored the potential of modifying the plant architecture of Jan by mutating the *ERECTA* (*ER*) gene (*StER*, Soltu.Jan_v1.1.08G009340.1). *ER* is known to control internode length in *A. thaliana* (Torii et al., 1996). Mutation of this gene in tomato, *SlER*, resulted in a compact and dwarf phenotype (Kwon et al., 2020). In order to target *StER* in Jan, we designed a 20-bp gRNA targeting exon 3 of *StER* (**Figure 6a**). We generated nine T0 plants with targeted edits in *StER* and five of these plants carried biallelic or homozygous mutations. We selected two of them for further analysis:

(1) *er-1*: both *StER* alleles were mutated in *er-1*, with 7 bp and 22 bp deletion, respectively. Both mutated alleles result in a shortened transcript caused by a premature stop codon. The two short transcripts would result in two truncated proteins that lose 886 aa and 891 aa, respectively (**Figure 6b**).
(2) *er-2*: a homozygous mutant, both *StER* alleles showed a 1-bp insertion, which encode a truncated protein that loses 885 aa (**Figure 6b**). The *er-2* mutant is named as mini-Jan^E^.

**Figure 6.**
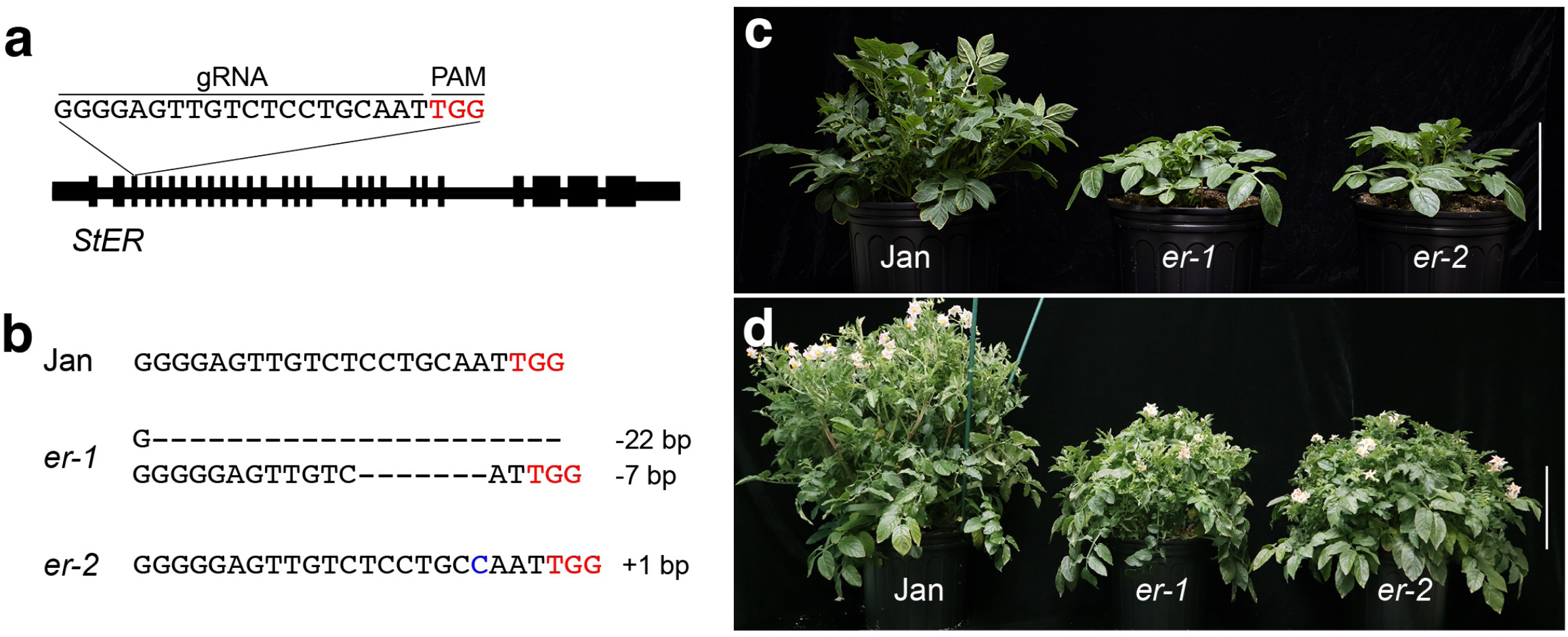
Genomic composition and phenotype of mini-Jan mutants from mutagenesis of the *StER* gene. (**a**) Diagram of the gRNA for CRISPR/Cas9 experiments targeting the *StER* gene. (**b**) Sequences of Jan, *er-1* and *er-2* in the genomic regions associated with mutations of the *StER* gene. (**c**) A single plant of Jan, *er-1* and *er-2* at 28 days after planting in a growth chamber. (**d)** A single plant of Jan, *er-1* and *er-2* at 48 days after planting in a growth chamber. All vertical bars = 20 cm.

Both *er-1* and *er-2* plants displayed a similar phenotype characterized by being shorter and more compact than the wild-type Jan **(Figure 6, c-d)**. Both mutants exhibited much tighter clustering of flowers, with shorter inflorescences compared to wild type **(Figure 6d)**. Both mutants also show reduced apical development, contributing to their more flattened architecture. The tissue culture seedlings of both *er-1* and *er-2* were especially pronounced for its short internode and dwarf phenotypes compared to the wild-type (**Figure 5b**). Interestingly, the *StER* mutants appear to have greater lateral growth, each possessing a thick central stem. This thick central stem is prominent in the mutants, with side branches comprising more than 90% of the plant, in contrast to the wild-type, which has many independent shoots and a less distinguishable central stem.

## Discussion

For some plant species, such as *A. thaliana*, transformation-mediated functional genomics studies can be performed universally for any genotype or accession. However, for a number of crop species, a model genotype or cultivar is required to be efficient for transformation-based research. For example, maize (*Zea mays*) was initially known to be recalcitrant to regeneration and transformation and most maize inbreds or hybrids are difficult to be transformed. Yet Hi-II (high type II callus production) maize has become the most extensively used maize line for transformation due to its exceptional ability to induce a high frequency of type II somatic embryogenic callus (Armstrong, 1999; Armstrong and Green, 1985). Unlike the more common and less regenerative type I callus, type II callus is friable and embryogenic (Yadava et al., 2017). The Hi-II line is a hybrid derived from two maize inbred lines, inbred A188 known for its favorable tissue culture characteristics (Lin et al., 2021) and inbred B73 for its superior agronomic qualities (Schnable et al., 2009). This combination has rendered the Hi-II line highly responsive in tissue culture and robust in field performance (Vega et al., 2008). Similarly, most varieties of common wheat (*Triticum aestivum*) are not amenable for transformation. Wheat laboratories have relied on a highly transformable variety “Bobwhite” (or its sister lines) for transgenic research (Pellegrineschi et al., 2002). In potato, although most tetraploid cultivars are amenable for transformation, tetraploid genotypes are not ideal for CRISPR/Cas-mediated gene editing.

The potato research community has long been searching for a model line for functional genomics studies. The DM potato line was chosen for genome sequencing largely due to its complete homozygosity (Pham et al., 2020; The Potato Genome Sequencing Consortium, 2011; Yang et al., 2023). Unfortunately, DM is weak, male sterile, and associated with poor tuber traits including the “jelly end” (tuber end rot) defect (Endelman and Jansky, 2016). Vigorous, highly fertile diploid potato lines with excellent tuber traits are available, such as RH (Zhou et al., 2020). However, most of the diploid lines are highly heterozygous and self-incompatible.

Although self-incompatible diploid lines can be converted to self-compatible by knocking out the *S-RNase* gene (Enciso-Rodriguez et al., 2019; Ye et al., 2018), selfed progenies from heterozygous diploids generally suffer from severe inbreeding depression. We demonstrate here that Jan has all the required characteristics as a model for functional genomics. Jan combines the most valuable traits from both parents: vigor, self-compatibility, and fertility from M6 and regeneration and transformation capability from DM. Jan shows good tuber traits under greenhouse condition. Jan tubers do not have the jelly end defect associated with DM and are considerably larger than those from M6. In addition, the compact and dwarf plant statue of mini-Jan will allow to grow more plants in limited greenhouse or growth chamber space.

A high or acceptable transformation efficiency is arguably the most important trait as a model for functional genomics. Jan showed a transformation efficiency of 10.2, 14.3, and 15.9% in three independent CRISPR/Cas9 experiments. The weighted average of these transformation efficiencies is 13.1%, which is comparable to the transformation efficiencies of other potato varieties. For example, when performing *Agrobacterium*-mediated transformation of stem internode explants of cultivars Lady Olympia, Granola, Agria, Désirée, and Innovator had transformation efficiencies of 22%, 20%, 18.6%, 15%, and 10%, respectively (Bakhsh, 2020).

Another example is AGB with a transformation efficiency of 11.5% and M6 having a transformation efficiency of 10% (Yasmeen et al., 2023). Comparing these varieties by ploidy level shows that tetraploids tend to have greater transformation efficiencies than diploids. This is further exemplified by diploid lines 3C (11.6%) and 10J (6.7%) having lower transformation efficiencies than tetraploid varieties (Nadolska-Orczyk et al., 2007). This phenomenon can be explained by treating T-DNA integration as a reaction and considering nuclear DNA as a substrate. Tetraploid varieties having approximately twice the amount of nuclear DNA than diploid varieties providing a plausible explanation for why tetraploids have higher transformation efficiencies.

It should be noted that the transformation efficiency of Jan can be improved in the future as our transformation procedure has not been optimized specifically for Jan. Nevertheless, the current transformation rate has already made Jan and mini-Jan a highly efficient system for gene editing due to its high-level homozygosity and two alleles for each gene. The technical challenge for editing all four alleles in tetraploid potato has been reported in several recent publications (Huang et al., 2023; Zhu et al., 2024). We were able to identify multiple biallelic or homozygous *StER* editing lines from merely nine transgenic Jan lines. We have achieved a similar editing rate of several other potato genes using Jan (unpublished) and were able to obtain biallelic or homozygous CRISPR/Cas lines in 4-6 months for the CRISPR projects. Consequenctly, gene editing experiments with Jan can be done as efficiently as in tomato. Thus, Jan and mini-Jan provide a highly efficient model system for gene editing and other transformation-mediated functional genomics studies.

## Experimental procedures

### Plant materials and growth conditions

The self-compatible diploid potato clone DMF5163 was derived from a cross between *S. tuberosum* Group Phureja DM 1-3 516 R44 (DM1-3) and *S. chacoense* (M6) (Endelman and Jansky, 2016). This clone has been self-pollinated for seven generations. Jan was propagated in vitro using nodal cuttings in tissue culture, grown on Murashige and Skoog (MS) medium (MS basal salts plus vitamins, 30 g/L sucrose, 4 g/L Gelrite, pH 5.8) (Murashige and Skoog, 1962). The plants were cultivated in culture tubes within growth chambers set to a 16-hour light/8-hour dark photoperiod at 22°C, with an average light intensity of 200 μmol m²s⁻¹.

Three-week-old plants were transplanted into a walk-in growth chamber with conditions of 16 hours of light at 22°C and 8 hours of darkness at 18°C, and a light intensity of 300 μmol m²s⁻¹ until flowering. Post-flowering, the plants were moved to a greenhouse for tuber collection. The greenhouse conditions were maintained at 16 hours of light at 24°C and 8 hours of darkness at 16°C, with a light intensity of 600 μmol m²s⁻¹, combining natural light with supplemental lighting from high-pressure sodium lamps.

### Pollen viability evaluation by I2-KI staining

Pollen was collected from a single flower of Jan and bulked. A 20 μL aliquot of I2-KI solution was mixed with the pollen and placed on a glass slide, then covered with a coverslip. Images of the pollen were captured using a QImaging Retiga EXi Fast 1394 CCD camera (Teledyne Photometrics, Tucson, AZ, USA) attached to an Olympus BX51 epifluorescence microscope. A field of view representative of the entire slide was used for analysis. Pollen grains that were stained yellow, round, and turgid were considered viable.

### Genome sequencing and assembly

Jan was grown in a growth chamber under 16h light at 22°C (8h dark at 18°C) and immature leaf tissue was harvested and flash-frozen in liquid nitrogen. High-molecular-weight genomic DNA was isolated via a crude Carlson lysis buffer extraction method and then cleaned with a Genomic-tip 500/G column (Qiagen, Hilden, Germany) to elute (Vaillancourt and Buell, 2019). Genomic DNA was size-selected using the Short Read Eliminator kit v1.0 (Circulomics, Pacific BioSciences, Menlo Park, CA); RNA was removed via digestion with RNase A (Qiagen) and subsequent re-purification of genomic DNA from the RNase A-digested solution. ONT sequencing libraries were prepared using the ONT SQK-LSK110 kit, loaded on R9.4.1 FLO-MIN106D flow cells, and sequenced by MinION Mk1B; the most recent ONT software available at the run dates of the sequencing libraries was used. The ONT whole-genome sequencing libraries were base-called using Guppy (v4.0.15, https://nanoporetech.com/community) using the high-accuracy model (dna_r9.4.1_450bps_hac), generating 65.5 Gb of sequencing data (**Table S1**). Reads shorter than 10 kb were filtered out using seqtk (v1.4-r130-dirty; https://github.com/lh3/seqtk), resulting in a final read set consisting of 1,927,473 reads amounting to 53.1 Gb (∼62.9x coverage). These reads were assembled using Flye (v.2.9.3-b1797; https://github.com/mikolmogorov/Flye) (Kolmogorov et al., 2019) with the parameters ‘--nano-raw’ and ‘--genome-size 0.8g’. Two iterations of error correction and polishing were performed with Racon (v1.5.0; https://github.com/lbcb-sci/racon) with the ‘-u’ parameter set; prior to each iteration of Racon, alignments of the final reads were generated in SAM format with Minimap2 (v2.26-r1175; https://github.com/lh3/minimap2) (Li, 2018) with the parameter ‘-ax map-ont’. The assembly was then polished by two rounds of Medaka (v1.11.3, https://github.com/nanoporetech/medaka) using the ‘r941_min_hac_g507’ model, followed by two rounds of NextPolish (v1.4.1, https://github.com/Nextomics/NextPolish) (Hu et al., 2020) using 55.7 Gb Illumina paired-end whole-genome shotgun reads (**Table S1**). Putative duplication within the assembly, indicated by evidence of residual heterozygosity from Illumina WGS sequencing data, as profiled by GenomeScope2.0 (https://github.com/tbenavi1/genomescope2.0), was removed with purge_dups (v.1.2.6, https://github.com/dfguan/purge_dups) (Guan et al., 2020) using default parameters and contigs shorter than 30 kb were filtered out with seqtk, producing a 717.2 Mb contig-level assembly consisting of 953 contigs with a contig N50 of 6.7 Mb (**Table S2**). The contigs were scaffolded with a reference-guided approach using RagTag ‘scaffold’ (v2.1.0, https://github.com/malonge/RagTag) (Alonge et al., 2022) with the parameters ‘-i 0.5’ and ‘-u’, and using the DM v6.1 genome assembly (Pham et al., 2020) as the reference, producing a 717.2 Mb chromosome-scale assembly composed of 84 total scaffolds with a scaffold N50 of 60.7 Mb (**Table S2**). Presence/absence of WGS k-mers in the Janv1.1 genome assembly was measured with K-mer Analysis Toolkit (KAT) (v2.4.2; https://github.com/TGAC/KAT) (Mapleson et al., 2017). Benchmarking Universal Single-Copy Orthologs (BUSCO) analysis (v5.4.3, https://busco.ezlab.org) (Manni et al., 2021; Simao et al., 2015) of the Jan assembly was performed using the embryophyta_odb10 lineage dataset and was run using Metaeuk (v7.bba0d80, https://github.com/soedinglab/metaeuk) (Karin et al., 2020) as the gene predictor in eukaryotic genome mode (**Table S3**).

### Genome annotation

A custom repeat library was generated from the contig-level assembly using RepeatModeler (v2.0.5; https://github.com/Dfam-consortium/RepeatModeler) (Flynn et al., 2020). The resulting repeat library was used to soft-mask the Jan genome assembly using RepeatMasker (v4.1.5; https://www.repeatmasker.org/RepeatMasker) (Tarailo-Graovac and Chen, 2009) with the following parameters: ‘-e ncbi -no_is -xsmall -gff (**Table S4**).

Empirical evidence for gene annotation included RNA-seq and full-length cDNA sequences. Jan was grown under 16h light at 22°C (8h dark at 18°C): young (immature) leaves and flower buds were collected from 6-week-old plants in the walk-in growth chamber; open (mature) flowers (6 weeks old), stolons, and young tubers (9 weeks old) were harvested from plants in the greenhouse; and roots were collected from 3-week-old tissue culture plantlets in the in-house growth chamber; all tissues were harvested by flash freezing in liquid nitrogen. RNA was extracted using the RNeasy Plant Mini Kit (Qiagen); on-column digestion with DNase I was performed. Stranded mRNA sequencing libraries were prepared using the KAPA mRNA HyperPrep Kit (Roche, Basel, Switzerland) and were sequenced on an S4 flow cell in paired-end mode on an Illumina NovaSeq 6000 (Illumina, San Diego, CA, USA), generating ∼40-50 million read pairs of length 150 nt per tissue type (**Table S1**). RNA-seq libraries were cleaned using Cutadapt (v4.6; (Martin, 2011)) with a minimum length of 100 nt and quality cutoff of 10. Cleaned reads were aligned to the repeat-masked genome using HISAT2 (v2.2.1; (Kim et al., 2019)) and alignment rates determined (**Table S1**). For generation of full-length cDNAs, the Dynabeads mRNA Purification Kit (ThermoFisher Scientific, Waltham, MA, Cat #61011) was used to isolate mRNA from the total RNA. cDNA libraries were constructed using Oxford Nanopore Technologies (ONT) SQK-PCS111 kit with the purified mRNA which were sequenced on FLO-MIN106 RevD flowcells using a MinION. Reads were base called using Dorado (v0.7.2; https://github.com/nanoporetech/dorado) with a minimum read mean quality score of 10, no trimming, using the model <u>dna_r9.4.1_e8_sup@v3.6</u> (**Table S1**). Pychopper (v2.5.0; https://github.com/nanoporetech/pychopper) was used to process the ONT full-length cDNA reads and trimmed reads greater than 500 nt were aligned to the Jan genome using Minimap2 (v2.17-r941; (Li, 2018)) with a maximum intron length of 5,000 nt. Aligned RNA-seq and ONT cDNA reads were assembled using Stringtie (v2.2.1; (Kovaka et al., 2019)); transcripts less then 500 nt were removed.

Using the soft-masked genome assemblies and empirical transcripts as hints, initial gene models were created using BRAKER2 (v2.1.6; (Bruna et al., 2021)). Initial gene models were then refined using two rounds of PASA2 (v2.5.2; (Campbell et al., 2006)) to create a working gene model set. High-confidence gene models were identified by filtering out gene models without expression evidence, or a PFAM domain match, or were a partial gene model or contained an interior stop codon. Functional annotation was assigned by searching the gene models proteins against the TAIR (v10; (Lamesch et al., 2012)) database and the Swiss-Prot plant proteins (release 2015_08) database using BLASTP (v2.12.0; (Altschul et al., 1990)) and the PFAM (v35.0; (El-Gebali et al., 2019)) database using PfamScan (v1.6; (Li et al., 2015)); functional descriptions were assigned based on the first significant hit. BUSCO analysis of the predicted protein sets produced from the annotation of the Jan assembly was performed using the embryophyta_odb10 lineage dataset and was run in proteins mode (**Table S3**).

### Synteny analysis

Genomic synteny between the Janv1.1, DMv6.1, and M6v5.0 genome assemblies was profiled and plotted on a chromosome-by-chromosome basis (with noise hidden) using D-GENIES (v1.5.0; https://dgenies.toulouse.inra.fr; (Cabanettes and Klopp, 2018)). GENESPACE (v1.3.1; (Lovell et al., 2022)) was used to identify syntelogs between Janv1.1 and other predicted proteomes and to construct riparian plots (**Figure S3**): DMv6.1 (Pham et al., 2020), M6v5.0 (http://spuddb.uga.edu/M6_v5_0_download.shtml), DM1S1 v1 (Jayakody et al., 2023), *S. candolleanum* v1.0 (http://spuddb.uga.edu/S_candolleanum_v1_0_download.shtml), and *S. lycopersicum* SL4.0 (Hosmani et al., 2019). Syntelogs between Janv1.1, DMv6.1, and M6v5.0 were identified by extracting all syntenic array members from the GENESPACE results using the query_pangenes() function (**Dataset 1**).

### Inherited genomic sequence assignment and GO enrichment of inherited genes

To identify DM and M6 genomic sequences inherited by Jan, k-mer indices were built for the Janv1.1, DMv6.1, and M6v5.0 genome assemblies; DM and M6 k-mers were anchored to the Jan assembly on a chromosome-by-chromosome basis and k-mer conservation between Jan and each parent was calculated in 100kb windows using PanKmer (v0.20.3; (Aylward et al., 2023)) with the parameter ‘--bin-size -1’. Windows displaying k-mer conservation differences of greater than 15% were assigned as inherited from the parent with the greater conservation level; having 15% or less k-mer conservation difference (i.e., high sequence conservation between DM and M6), the remaining windows were designated as being ambiguous. These categorized genomic regions were plotted using karyoploteR (v1.30.0; https://bioconductor.org/packages/release/bioc/html/karyoploteR.html) (Gel and Serra, 2017). Genes were assigned as being inherited from DM, M6, or as being of ambiguous inheritance based on their positions within these calculated genomic sequence blocks using BEDTools (v2.31.1; https://github.com/arq5x/bedtools2) (Quinlan and Hall, 2010).

For biological analysis of the inherited genes, Gene Ontology (GO) terms were assigned to Jan genes using InterProScan (v5.63-95.0; https://github.com/ebi-pf-team/interproscan, (Blum et al., 2021; Jones et al., 2014)). GO term enrichments for biological processes and molecular functions were performed on the groups of genes inherited from DM and from M6 using topGO (v2.54.0; https://bioconductor.org/packages/release/bioc/html/topGO.html) (Alexa et al., 2006) with the “classic” algorithm, and Fisher’s exact test was used for statistically significant enrichment. GO terms deemed significantly enriched (p-value < 0.01) were extracted and summarized with Revigo (v1.8.1; http://revigo.irb.hr) (Supek et al., 2011) using the whole UniProt database (**Table S5**). To incorporate expression evidence into this biological analysis, gene expression levels for each tissue type in Jan were quantified for the representative high-confidence gene models in transcripts per million (TPM) from the trimmed RNA-seq reads using Kallisto (v0.50.1; https://github.com/pachterlab/kallisto) (Bray et al., 2016) with the parameter ‘--rf-stranded’ (**Dataset 2**).

### CRISPR–Cas9 vector construction

The specific gRNAs used in this study were designed using CRISPR RGEN tools (http://www.rgenome.net/cas-designer/). The CRISPR/Cas9 mutagenesis vectors were generated following published protocols (Cermak et al., 2017). Briefly, the gRNA was cloned into the pMOD_B2515 vector using a Golden Gate reaction with Esp3I to create the *AtU6::gRNA* cassette. Subsequently, the *AtU6::gRNA* and Cas9 expression cassette (*CmYLCV::Cas9*) was assembled into the binary vectors pTRANS_220d or pTRANS_210d using a Golden Gate reaction with AarI, resulting in the final CRISPR/Cas9 mutagenesis vector.

### *Agrobacterium*-mediated transformation

CRISPR-Cas9 constructs were transformed into *A. tumefaciens* GV3101 (pMP90) and incubated on LB agar containing 50 mg/L Gentamycin and 50 mg/L Kanamycin. The *Agrobacterium*-mediated transformation of Jan was performed using internode explants, following previously published protocols with some modifications (Nadakuduti et al., 2019; Paz and Veilleux, 1999). Internodes were excised from four-week-old healthy in vitro plants and placed horizontally on pre-culture media (MS basal salts plus vitamins, 30 g/L sucrose, 8 g/L agar, 2 mg/L 2,4-D, 0.8 mg/L Zeatin Riboside) for two days.

*Agrobacterium* seed cultures were prepared by inoculating liquid LB with a single positive colony, followed by overnight incubation at 28°C with shaking at 220 rpm. The next day, liquid cultures were diluted 1:50 in LB and continued shaking until reaching an OD_600_ of 0.6. The internodes were then incubated in the *Agrobacterium* culture suspended in MS liquid media (MS basal salts plus vitamins, 30 g/L sucrose) for 15 minutes. Post-infection, the internodes were dried on sterilized filter paper and placed onto co-culture medium (same composition as pre-culture medium) with a piece of filter paper for three days in the dark.

After three days of co-cultivation, the internodes were washed six times with sterile double-distilled water (ddH_2_O) containing 400 mg/L Timentin. The dried internodes were then transferred to callus-induction medium (MS basal salts plus vitamins, 30 g/L sucrose, 8 g/L agar, 2.5 mg/L Zeatin Riboside, 0.1 mg/L NAA, 0.2 mg/L GA3, 400 mg/L Timentin, and either 50 mg/L Kanamycin or 5 mg/L Hygromycin) for one week. Subsequently, the internodes were transferred to shoot-induction medium (MS basal salts plus vitamins, 30 g/L sucrose, 8 g/L agar, 1 mg/L Zeatin Riboside, 2 mg/L GA3, 400 mg/L Timentin, and either 100 mg/L Kanamycin or 10 mg/L Hygromycin). Explants were transferred to fresh shoot-induction medium weekly until shoots emerged. Once shoots emerged, the explants were moved to shoot-elongation medium (MS basal salts plus vitamins, 30 g/L sucrose, 8 g/L agar, 400 mg/L Timentin, and either 100 mg/L Kanamycin or 10 mg/L Hygromycin).When the shoots reached 1–2 cm in length, they were gently cut slightly above the base and transferred to root-induction medium (MS basal salts plus vitamins, 30 g/L sucrose, 8 g/L agar, 200 mg/L Timentin, and either 50 mg/L Kanamycin or 5 mg/L Hygromycin).

### Detection of targeted mutations

DNA was extracted from the young leaves of the rooted plants using the DNeasy Plant Mini Kit (Qiagen, Hilden, Germany). Positive T0 plants were screened by PCR using transgene-specific primers. To detect mutations at the target site, PCR for target site amplification was performed using specific primers, followed by direct sequencing with Sanger sequencing technology. The Sanger sequencing results were analyzed using ICE software (Conant et al., 2022) and TIDE software (Brinkman et al., 2014) to determine the mutation types.

## Supporting information

**Figure S1.** Pollen and seeds of Jan.

**Figure S2.** Heterozygous residue in the Jan genome.

**Figure S3.** Riparian plot displaying genic synteny between Jan and other diploid Solanum species.

**Figure S4.** Pairwise collinearity analysis between Jan and both DM and M6.

**Figure S5.** Allelic representation of DMv6.1 and M6v5.0 genomes in the Janv1.1 genome assembly.

**Table S1.** DNA and RNA sequencing information for Jan.

**Table S2.** Genome assembly summary and statistics of Jan.

**Table S3.** BUSCO scores for Jan genome assembly and annotation.

**Table S4.** Repetitive DNA in the Jan genome.

**Table S5.** GO terms enriched in Jan genes inherited from DM and M6.

**Dataset 1.** Janv1.1 genes and their syntelogs in DMv6.1 and M6v5.0.

**Dataset 2.** Expression of Jan genes, measured in transcripts per million.

## Acknowledgments

We thank Brieanne Kniahynycky for her assistance in genomic sequence data management and thank Brieanne Kniahynycky and Joshua Wood for their assistance with full-length cDNA sequencing. The research described in this study was supported by the National Institute of General Medical Sciences of the National Institutes of Health under Award Numbers T32GM110523 and T32GM152798 to L.W.S. and J.K.T; by funds from the Georgia Research Alliance, Georgia Seed Development, and the University of Georgia to C.R.B.; by grants IS-5317-20C and IS-5684-24C from BARD (the United States - Israel Binational Agricultural Research and Development Fund), AgBioResearch at Michigan State University (Hatch grant MICL02571), and MSU startup funds to J.J.

## Data and material availability

All sequencing reads are available in the National Center for Biotechnology Information Sequence Read Archive under BioProject PRJNA1157315. The genome sequence of Jan is downloadable (https://spuddb.uga.edu/jan_v1_1_download.shtml) and the annotation can be viewed at SpudDB (https://spuddb.uga.edu/download.shtml). Seeds from Jan and mini-Jan are available upon request. Seeds will be sent after the requestors complete the relevant Plant Quarantine forms from the requestor’s country.

## Author contributions

J.J. conceived the research. H.X., L.W.S., J.K.T., and N.M.B. conducted the experiments. J.P.H., C.F., D.S.D., C.R.B., and J.J. analyzed the data. H.X., L.W.S., J.K.T., C.B.R., and J.J. wrote the manuscript.

## References

Achakkagari, S.R., Kyriakidou, M., Gardner, K.M., De Koeyer, D., De Jong, H., Strömvik, M. and Tai, H.H. (2022) Genome sequencing of adapted diploid potato clones. Frontiers in Plant Science 13, 954933.

Alexa, A., Rahnenführer, J. and Lengauer, T. (2006) Improved scoring of functional groups from gene expression data by decorrelating GO graph structure. Bioinformatics 22, 1600–1607.

Alonge, M., Lebeigle, L., Kirsche, M., Jenike, K., Ou, S.J., Aganezov, S., Wang, X.A., Lippman, Z.B., Schatz, M.C. and Soyk, S. (2022) Automated assembly scaffolding using RagTag elevates a new tomato system for high-throughput genome editing. Genome Biol 23, 258.

Alsahlany, M., Enciso-Rodriguez, F., Lopez-Cruz, M., Coombs, J. and Douches, D.S. (2021) Developing self-compatible diploid potato germplasm through recurrent selection. Euphytica 217, 47.

Altschul, S.F., Gish, W., Miller, W., Myers, E.W. and Lipman, D.J. (1990) Basic local alignment search tool. Journal of Molecular Biology 215, 403–410.

Armstrong, C.L. (1999) The first decade of maize transformation: A review and future perspective. Maydica 44, 101–109.

Armstrong, C.L. and Green, C.E. (1985) Establishment and maintenance of friable, embryogenic maize callus and the involvement of L-proline. Planta 164, 207–214.

Aylward, A.J., Petrus, S., Mamerto, A., Hartwick, N.T. and Michael, T.P. (2023) PanKmer: k-mer-based and reference-free pangenome analysis. Bioinformatics 39, btad621.

Bais, H.P., Sudha, G.S. and Ravishankar, G.A. (2000) Putrescine and silver nitrate influences shoot multiplication, in vitro flowering and endogenous titers of polyamines in *Cichorium intybus* L. cv. Lucknow local. Journal of Plant Growth Regulation 19, 238–248.

Bakhsh, A. (2020) Development of efficient, reproducible and stable Agrobacterium-mediated genetic transformation of five potato cultivars. Food Technol Biotech 58, 57–63.

Ballvora, A., Ercolano, M.R., Weiss, J., Meksem, K., Bormann, C.A., Oberhagemann, P., Salamini, F. and Gebhardt, C. (2002) The *R1* gene for potato resistance to late blight (*Phytophthora infestans*) belongs to the leucine zipper/NBS/LRR class of plant resistance genes. Plant J. 30, 361–371.

Bethke, P.C., Halterman, D.A., Francis, D.M., Jiang, J.M., Douches, D.S., Charkowski, A.O. and Parsons, J. (2022) Diploid potatoes as a catalyst for change in the potato industry. American Journal of Potato Research 99, 337–357.

Bishop, G.J., Nomura, T., Yokota, T., Harrison, K., Noguchi, T., Fujioka, S., Takatsuto, S., Jones, J.D.G. and Kamiya, Y. (1999) The tomato DWARF enzyme catalyses C-6 oxidation in brassinosteroid biosynthesis. P Natl Acad Sci USA 96, 1761–1766.

Blum, M., Chang, H.Y., Chuguransky, S., Grego, T., Kandasaamy, S., Mitchell, A., Nuka, G., Paysan-Lafosse, T., Qureshi, M., Raj, S., Richardson, L., Salazar, G.A., Williams, L., Bork, P., Bridge, A., Gough, J., Haft, D.H., Letunic, I., Marchler-Bauer, A., Mi, H.Y., Natale, D.A., Necci, M., Orengo, C.A., Pandurangan, A.P., Rivoire, C., Sigrist, C.J.A., Sillitoe, I., Thanki, N., Thomas, P.D., Tosatto, S.C.E., Wu, C.H., Bateman, A. and Finn, R.D. (2021) The InterPro protein families and domains database: 20 years on. Nucleic Acids Research 49, D344–D354.

Bonierbale, M.W., Plaisted, R.L. and Tanksley, S.D. (1988) RFLP maps based on a common set of clones reveal modes of chromosomal evolution in potato and tomato. Genetics 120, 1095–1103.

Bray, N.L., Pimentel, H., Melsted, P. and Pachter, L. (2016) Near-optimal probabilistic RNA-seq quantification. Nature Biotechnology 34, 525–527.

Brinkman, E.K., Chen, T., Amendola, M. and van Steensel, B. (2014) Easy quantitative assessment of genome editing by sequence trace decomposition. Nucleic Acids Research 42, e168.

Bruna, T., Hoff, K.J., Lomsadze, A., Stanke, M. and Borodovsky, M. (2021) BRAKER2: automatic eukaryotic genome annotation with GeneMark-EP plus and AUGUSTUS supported by a protein database. Nar Genomics and Bioinformatics 3, lqaa108.

Burtin, D., Martintanguy, J., Paynot, M. and Rossin, N. (1989) Effects of the suicide inhibitors of arginine and ornithine decarboxylase activities on organogenesis, growth, free polyamine and hydroxycinnamoyl putrescine levels in leaf explants of Nicotiana Xanthi n.c. cultivated in vitro in a medium producing callus formation. Plant Physiology 89, 104–110.

Cabanettes, F. and Klopp, C. (2018) D-GENIES: dot plot large genomes in an interactive, efficient and simple way. Peerj 6, e4958.

Campbell, M.A., Haas, B.J., Hamilton, J.P., Mount, S.M. and Buell, C.R. (2006) Comprehensive analysis of alternative splicing in rice and comparative analyses with Arabidopsis. Bmc Genomics 7, 327.

Cermak, T., Curtin, S.J., Gil-Humanes, J., Cegan, R., Kono, T.J.Y., Konecna, E., Belanto, J.J., Starker, C.G., Mathre, J.W., Greenstein, R.L. and Voytasa, D.F. (2017) A multipurpose toolkit to enable advanced genome engineering in plants. Plant Cell 29, 1196–1217.

Conant, D., Hsiau, T., Rossi, N., Oki, J., Maures, T., Waite, K., Yang, J.Y., Joshi, S., Kelso, R., Holden, K., Enzmann, B.L. and Stoner, R. (2022) Inference of CRISPR edits from Sanger trace data. Crispr J 5, 123–130.

Crane, Y.M. and Gelvin, S.B. (2007) RNAi-mediated gene silencing reveals involvement of chromatin-related genes in *Agrobacterium*-mediated root transformation. P Natl Acad Sci USA 104, 15156–15161.

de Vries, M.E., Adams, J.R., Eggers, E.J., Ying, S., Stockem, J.E., Kacheyo, O.C., van Dijk, L.C.M., Khera, P., Bachem, C.W., Lindhout, P. and van der Vossen, E.A.G. (2023) Converting hybrid potato breeding science into practice. Plants-Basel 12, 230.

Devaux, A., Goffart, J.-P., Petsakos, A., Kromann, P., Gatto, M., Okello, J., Suarez, V. and Hareau, G. (2020) Global food security, contributions from sustainable potato agri-food systems. In: The potato crop: its agricultural, nutritional and social contribution to humankind (Campos, H., Ortiz, O. ed) pp. 3–35. Dordrecht: Springer.

Douches, D.S., Maas, D., Jastrzebski, K. and Chase, R.W. (1996) Assessment of potato breeding progress in the USA over the last century. Crop Sci. 36, 1544–1552.

Duangpan, S., Zhang, W.L., Wu, Y.F., Jansky, S.H. and Jiang, J.M. (2013) Insertional mutagenesis using *Tnt1* retrotransposon in potato. Plant Physiology 163, 21–29.

Eggers, E.J., van der Burgt, A., van Heusden, S.A.W., de Vries, M.E., Visser, R.G.F., Bachem, C.W.B. and Lindhout, P. (2021) Neofunctionalisation of the *Sli* gene leads to self-compatibility and facilitates precision breeding in potato. Nature Communications 12, 4141.

El-Gebali, S., Mistry, J., Bateman, A., Eddy, S.R., Luciani, A., Potter, S.C., Qureshi, M., Richardson, L.J., Salazar, G.A., Smart, A., Sonnhammer, E.L.L., Hirsh, L., Paladin, L., Piovesan, D., Tosatto, S.C.E. and Finn, R.D. (2019) The Pfam protein families database in 2019. Nucleic Acids Research 47, D427–D432.

Enciso-Rodriguez, F., Manrique-Carpintero, N.C., Nadakuduti, S.S., Buell, C.R., Zarka, D. and Douches, D. (2019) Overcoming self-incompatibility in diploid potato using CRISPR-Cas9. Frontiers in Plant Science 10, 376.

Endelman, J.B. and Jansky, S.H. (2016) Genetic mapping with an inbred line-derived F2 population in potato. Theor Appl Genet 129, 935–943.

Fazilati, M. and Forghani, A.H. (2015) The role of polyamine to increasing growth of plant: as a key factor in health crisis. Int. J. Health Syst. Disaster Manag. 3, 89–94.

Flynn, J.M., Hubley, R., Goubert, C., Rosen, J., Clark, A.G., Feschotte, C. and Smit, A.F. (2020) RepeatModeler2 for automated genomic discovery of transposable element families. P Natl Acad Sci USA 117, 9451–9457.

Gebhardt, C., Ritter, E., Debener, T., Schachtschabel, U., Walkemeier, B., Uhrig, H. and Salamini, F. (1989) RFLP analysis and linkage mapping in *Solanum tuberosum*. Theor. Appl. Genet. 78, 65–75.

Gel, B. and Serra, E. (2017) karyoploteR: an R/Bioconductor package to plot customizable genomes displaying arbitrary data. Bioinformatics 33, 3088–3090.

Guan, D.F., McCarthy, S.A., Wood, J., Howe, K., Wang, Y.D. and Durbin, R. (2020) Identifying and removing haplotypic duplication in primary genome assemblies. Bioinformatics 36, 2896–2898.

Guo, D., Wong, W.S., Xu, W.Z., Sun, F.F., Qing, D.J. and Li, N. (2011) *Cis-cinnamic acid-enhanced 1* gene plays a role in regulation of Arabidopsis bolting. Plant Molecular Biology 75, 481–495.

Hao, Y.J., Kitashiba, H., Honda, C., Nada, K. and Moriguchi, T. (2005) Expression of arginine decarboxylase and ornithine decarboxylase genes in apple cells and stressed shoots. J Exp Bot 56, 1105–1115.

Hoopes, G., Meng, X.X., Hamilton, J.P., Achakkagari, S.R., Guesdes, F.D.F. et al. (2022) Phased, chromosome-scale genome assemblies of tetraploid potato reveal a complex genome, transcriptome, and predicted proteome landscape underpinning genetic diversity. Mol Plant 15, 520–536.

Hosaka, A.J., Sanetomo, R. and Hosaka, K. (2022) A de novo genome assembly of *Solanum verrucosum* Schlechtendal, a Mexican diploid species geographically isolated from other diploid A-genome species of potato relatives. G3-Genes Genomes Genetics 12, jkac166.

Hosaka, K. and Sanetomo, R. (2020) Creation of a highly homozygous diploid potato using the locus inhibitor (*Sli*) gene. Euphytica 216, 169.

Hosmani, P.S., Flores-Gonzalez, M., van de Geest, H., Maumus, F., Bakker, L.V., Schijlen, E., van Haarst, J., Cordewener, J., Sanchez-Perez, G., Peters, S., Fei, Z., Giovannoni, J.J., Mueller, L.A. and Saha, S. (2019) An improved de novo assembly and annotation of the tomato reference genome using single-molecule sequencing, Hi-C proximity ligation and optical maps. bioRxiv, 767764.

Hu, J., Fan, J.P., Sun, Z.Y. and Liu, S.L. (2020) NextPolish: a fast and efficient genome polishing tool for long-read assembly. Bioinformatics 36, 2253–2255.

Huang, X.E., Jia, H.G., Xu, J., Wang, Y.C., Wen, J.W. and Wang, N. (2023) Transgene-free genome editing of vegetatively propagated and perennial plant species in the T0 generation via a co-editing strategy. Nature Plants 9, 1591–1597.

Jansky, S.H., Charkowski, A.O., Douches, D.S., Gusmini, G., Richael, C., Bethke, P.C., Spooner, D.M., Novy, R.G., De Jong, H., De Jong, W.S., Bamberg, J.B., Thompson, A.L., Bizimungu, B., Holm, D.G., Brown, C.R., Haynes, K.G., Sathuvalli, V.R., Veilleux, R.E., Miller, J.C., Bradeen, J.M. and Jiang, J.M. (2016) Reinventing potato as a diploid inbred line-based crop. Crop Science 56, 1412–1422.

Jansky, S.H., Chung, Y.S. and Kittipadukal, P. (2014) M6: a diploid potato inbred line for use in breeding and genetics research. Journal of Plant Registrations 8, 195–199.

Jayakody, T.B., Enciso-Rodríguez, F.E., Jensen, J., Douches, D.S. and Nadakuduti, S.S. (2022) Evaluation of diploid potato germplasm for applications of genome editing and genetic engineering. American Journal of Potato Research 99, 13–24.

Jayakody, T.B., Hamilton, J.P., Jensen, J., Sikora, S., Wood, J.C., Douches, D.S. and Buell, C.R. (2023) Genome Report: Genome sequence of 1S1, a transformable and highly regenerable diploid potato for use as a model for gene editing and genetic engineering. G3-Genes Genomes Genetics 13, jkad036.

Jones, P., Binns, D., Chang, H.Y., Fraser, M., Li, W.Z., McAnulla, C., McWilliam, H., Maslen, J., Mitchell, A., Nuka, G., Pesseat, S., Quinn, A.F., Sangrador-Vegas, A., Scheremetjew, M., Yong, S.Y., Lopez, R. and Hunter, S. (2014) InterProScan 5: genome-scale protein function classification. Bioinformatics 30, 1236–1240.

Karin, E.L., Mirdita, M. and Söding, J. (2020) MetaEuk-sensitive, high-throughput gene discovery, and annotation for large-scale eukaryotic metagenomics. Microbiome 8.

Kim, D., Paggi, J.M., Park, C., Bennett, C. and Salzberg, S.L. (2019) Graph-based genome alignment and genotyping with HISAT2 and HISAT-genotype. Nature Biotechnology 37, 907–915.

Kim, H.J., Ok, S.H., Bahn, S.C., Jang, J., Oh, S.A., Park, S.K., Twell, D., Ryu, S.B. and Shin, J.S. (2011) Endoplasmic reticulum– and Golgi-localized phospholipase A2 plays critical roles in Arabidopsis pollen development and germination. Plant Cell 23, 94–110.

Kloosterman, B., Abelenda, J.A., Gomez, M.D.C., Oortwijn, M., de Boer, J.M., Kowitwanich, K., Horvath, B.M., van Eck, H.J., Smaczniak, C., Prat, S., Visser, R.G.F. and Bachem, C.W.B. (2013) Naturally occurring allele diversity allows potato cultivation in northern latitudes. Nature 495, 246–250.

Kolmogorov, M., Yuan, J., Lin, Y. and Pevzner, P.A. (2019) Assembly of long, error-prone reads using repeat graphs. Nature Biotechnology 37, 540–546.

Kovaka, S., Zimin, A.V., Pertea, G.M., Razaghi, R., Salzberg, S.L. and Pertea, M. (2019) Transcriptome assembly from long-read RNA-seq alignments with StringTie2. Genome Biol 20, 278.

Kwon, C.T., Heo, J., Lemmon, Z.H., Capua, Y., Hutton, S.F., Van Eck, J., Park, S.J. and Lippman, Z.B. (2020) Rapid customization of Solanaceae fruit crops for urban agriculture. Nature Biotechnology 38, 182–188.

Lamesch, P., Berardini, T.Z., Li, D.H., Swarbreck, D., Wilks, C., Sasidharan, R., Muller, R., Dreher, K., Alexander, D.L., Garcia-Hernandez, M., Karthikeyan, A.S., Lee, C.H., Nelson, W.D., Ploetz, L., Singh, S., Wensel, A. and Huala, E. (2012) The Arabidopsis Information Resource (TAIR): improved gene annotation and new tools. Nucleic Acids Research 40, D1202–D1210.

Leisner, C.P., Hamilton, J.P., Crisovan, E., Manrique-Carpintero, N.C., Marand, A.P., Newton, L., Pham, G.M., Jiang, J.M., Douches, D.S., Jansky, S.H. and Buell, C.R. (2018) Genome sequence of M6, a diploid inbred clone of the high-glycoalkaloid-producing tuber-bearing potato species *Solanum chacoense*, reveals residual heterozygosity. Plant Journal 94, 562–570.

Li, H. (2018) Minimap2: pairwise alignment for nucleotide sequences. Bioinformatics 34, 3094–3100.

Li, W.Z., Cowley, A., Uludag, M., Gur, T., McWilliam, H., Squizzato, S., Park, Y.M., Buso, N. and Lopez, R. (2015) The EMBL-EBI bioinformatics web and programmatic tools framework. Nucleic Acids Research 43, W580–W584.

Lin, G.F., He, C., Zheng, J., Koo, D.H., Le, H., Zheng, H.K., Tamang, T.M., Lin, J.G., Liu, Y., Zhao, M.X., Hao, Y.F., McFraland, F., Wang, B., Qin, Y., Tang, H.B., McCarty, D.R., Wei, H.R., Cho, M.J., Park, S., Kaeppler, H., Kaeppler, S.M., Liu, Y.J., Springer, N., Schnable, P.S., Wang, G.Y., White, F.F. and Liu, S.Z. (2021) Chromosome-level genome assembly of a regenerable maize inbred line A188. Genome Biol 22, 175.

Lovell, J.T., Sreedasyam, A., Schranz, M.E., Wilson, M., Carlson, J.W., Harkess, A., Emms, D., Goodstein, D.M. and Schmutz, J. (2022) GENESPACE tracks regions of interest and gene copy number variation across multiple genomes. Elife 11, e78526.

Ma, L., Zhang, C.Z., Zhang, B., Tang, F., Li, F.T., Liao, Q.G., Tang, D., Peng, Z., Jia, Y.X., Gao, M., Guo, H., Zhang, J.Z., Luo, X.M., Yang, H.Q., Gao, D.L., Lucas, W.J., Li, C.H., Huang, S.W. and Shang, Y. (2021) A *nonS-locus F-box* gene breaks self-incompatibility in diploid potatoes. Nature Communications 12, 4142.

Manni, M., Berkeley, M.R., Seppey, M., Simao, F.A. and Zdobnov, E.M. (2021) BUSCO update: novel and streamlined workflows along with broader and deeper phylogenetic coverage for scoring of eukaryotic, prokaryotic, and viral genomes. Mol Biol Evol 38, 4647–4654.

Mapleson, D., Accinelli, G.G., Kettleborough, G., Wright, J. and Clavijo, B.J. (2017) KAT: a K-mer analysis toolkit to quality control NGS datasets and genome assemblies. Bioinformatics 33, 574–576.

Mari, R.S., Schrinner, S., Finkers, R., Ziegler, F.M.R., Arens, P., Schmidt, M.H.W., Usadel, B., Klau, G.W. and Marschall, T. (2024) Haplotype-resolved assembly of a tetraploid potato genome using long reads and low-depth offspring data. Genome Biol 25, 26.

Marti, E., Gisbert, C., Bishop, G.J., Dixon, M.S. and Garcia-Martinez, J.L. (2006) Genetic and physiological characterization of tomato cv. Micro-Tom. J Exp Bot 57, 2037–2047.

Martin, M. (2011) Cutadapt removes adapter sequences from high-throughput sequencing reads. 2011 17, 10–12.

Martin-Tanguy, J. (2001) Metabolism and function of polyamines in plants: recent development (new approaches). Plant Growth Regulation 34, 135–148.

Murashige, T. and Skoog, F. (1962) A revised medium for rapid growth and bio assays with tobacco tissue cultures. Physiologia Plantarum 15, 473–497.

Mysore, K.S., Nam, J. and Gelvin, S.B. (2000) An Arabidopsis histone H2A mutant is deficient in Agrobacterium T-DNA integration. P Natl Acad Sci USA 97, 948–953.

Nadakuduti, S.S., Starker, C.G., Voytas, D.F., Buell, C.R. and Douches, D.S. (2019) Genome editing in potato with CRISPR/Cas9. In: Plant Genome Editing with Crispr Systems: Methods and Protocols (Qi, Y. ed) pp. 183–201.

Nadolska-Orczyk, A., Pietrusinska, A., Binka-Wyrwa, A., Kuc, D. and Orczyk, W. (2007) Diploid potato (L.) as a model crop to study transgene expression. Cell Mol Biol Lett 12, 206–219.

Ou, S.J., Chen, J.F. and Jiang, N. (2018) Assessing genome assembly quality using the LTR Assembly Index (LAI). Nucleic Acids Research 46, e126.

Paz, M.M. and Veilleux, R.E. (1999) Influence of culture medium and *in vitro* conditions on shoot regeneration in *Solanum phureja* monoploids and fertility of regenerated doubled monoploids. Plant Breeding 118, 53–57.

Pedersen, J.F., Bean, S.R., Funnell, D.L. and Graybosch, R.A. (2004) Rapid iodine staining techniques for identifying the waxy phenotype in sorghum grain and waxy genotype in sorghum pollen. Crop Science 44, 764–767.

Pellegrineschi, A., Noguera, L.M., Skovmand, B., Brito, R.M., Velazquez, L., Salgado, M.M., Hernandez, R., Warburton, M. and Hoisington, D. (2002) Identification of highly transformable wheat genotypes for mass production of fertile transgenic plants. Genome 45, 421–430.

Peng, X.Q., Wang, M.L., Li, Y.Q., Yan, W., Chang, Z.Y., Chen, Z.F., Xu, C.J., Yang, C.W., Deng, X.W., Wu, J.X. and Tang, X.Y. (2020) Lectin receptor kinase OsLecRK-S.7 is required for pollen development and male fertility. Journal of Integrative Plant Biology 62, 1227–1245.

Pham, G.M., Hamilton, J.P., Wood, J.C., Burke, J.T., Zhao, H.N., Vaillancourt, B., Ou, S.J., Jiang, J.M. and Buell, C.R. (2020) Construction of a chromosome-scale long-read reference genome assembly for potato. Gigascience 9, giaa100.

Quinlan, A.R. and Hall, I.M. (2010) BEDTools: a flexible suite of utilities for comparing genomic features. Bioinformatics 26, 841–842.

Ranallo-Benavidez, T.R., Jaron, K.S. and Schatz, M.C. (2020) GenomeScope 2.0 and Smudgeplot for reference-free profiling of polyploid genomes. Nature Communications 11, 1432.

Schnable, P.S., Ware, D., Fulton, R.S., Stein, J.C., Wei, F.S. et al. (2009) The B73 maize genome: Complexity, diversity, and dynamics. Science 326, 1112–1115.

Siegfried, K.R., Eshed, Y., Baum, S.F., Otsuga, D., Drews, G.N. and Bowman, J.L. (1999) Members of the *YABBY* gene family specify abaxial cell fate in *Arabidopsis*. Development 126, 4117–4128.

Simao, F.A., Waterhouse, R.M., Ioannidis, P., Kriventseva, E.V. and Zdobnov, E.M. (2015) BUSCO: assessing genome assembly and annotation completeness with single-copy orthologs. Bioinformatics 31, 3210–3212.

Song, J.Q., Bradeen, J.M., Naess, S.K., Raasch, J.A., Wielgus, S.M., Haberlach, G.T., Liu, J., Kuang, H.H., Austin-Phillips, S., Buell, C.R., Helgeson, J.P. and Jiang, J.M. (2003) Gene *RB* cloned from *Solanum bulbocastanum* confers broad spectrum resistance to potato late blight. Proc. Natl. Acad. Sci. U. S. A. 100, 9128–9133.

Stavolone, L., Kononova, M., Pauli, S., Ragozzino, A., de Haan, P., Milligan, S., Lawton, K. and Hohn, T. (2003) Cestrum yellow leaf curling virus (CmYLCV) promoter: a new strong constitutive promoter for heterologous gene expression in a wide variety of crops. Plant Molecular Biology 53, 703–713.

Sun, H.Q., Jiao, W.B., Campoy, J.A., Krause, K., Goel, M., Folz-Donahue, K., Kukat, C., Huettel, B. and Schneeberger, K. (2022) Chromosome-scale and haplotype-resolved genome assembly of a tetraploid potato cultivar. Nat Genet 54, 342–348.

Supek, F., Bosnjak, M., Skunca, N. and Smuc, T. (2011) REVIGO summarizes and visualizes long lists of gene ontology terms. Plos One 6, e21800.

Tarailo-Graovac, M. and Chen, N. (2009) Using RepeatMasker to identify repetitive elements in genomic sequences. Curr Protoc Bioinformatics Chapter 4, 4.10.11–14.10.14.

The Potato Genome Sequencing Consortium (2011) Genome sequence and analysis of the tuber crop potato. Nature 475, 189–195.

Torii, K.U., Mitsukawa, N., Oosumi, T., Matsuura, Y., Yokoyama, R., Whittier, R.F. and Komeda, Y. (1996) The Arabidopsis *ERECTA* gene encodes a putative receptor protein kinase with extracellular leucine-rich repeats. Plant Cell 8, 735–746.

Vaillancourt, B. and Buell, C.R. (2019) High molecular weight DNA isolation method from diverse plant species for use with Oxford Nanopore sequencing. bioRxiv, 783159.

Vega, J.M., Yu, W.C., Kennon, A.R., Chen, X.L. and Zhang, Z.Y.J. (2008) Improvement of *Agrobacterium*-mediated transformation in Hi-II maize (*Zea mays*) using standard binary vectors. Plant Cell Reports 27, 297–305.

Wan, J.R., Patel, A., Mathieu, M., Kim, S.Y., Xu, D. and Stacey, G. (2008) A lectin receptor-like kinase is required for pollen development in Arabidopsis. Plant Molecular Biology 67, 469–482.

Yadava, P., Abhishek, A., Singh, R., Singh, I., Kaul, T., Pattanayak, A. and Agrawal, P.K. (2017) Advances in maize transformation technologies and development of transgenic maize. Frontiers in Plant Science 7, 1949.

Yang, X.H., Zhang, L.K., Guo, X., Xu, J.F., Zhang, K., Yang, Y.Q., Yang, Y., Jian, Y.Q., Dong, D.F., Huang, S.W., Cheng, F. and Li, G.C. (2023) The gap-free potato genome assembly reveals large tandem gene clusters of agronomical importance in highly repeated genomic regions. Mol Plant 16, 314–317.

Yasmeen, A., Bakhsh, A., Ajmal, S., Muhammad, M., Sadaqat, S., Awais, M., Azam, S., Latif, A., Shahid, N. and Rao, A.Q. (2023) CRISPR/Cas9-mediated genome editing in diploid and tetraploid potatoes. Acta Physiol Plant 45, 32.

Ye, M.W., Peng, Z., Tang, D., Yang, Z.M., Li, D.W., Xu, Y.M., Zhang, C.Z. and Huang, S.W. (2018) Generation of self-compatible diploid potato by knockout of *S-RNase*. Nature Plants 4, 651–654.

Yi, H., Sardesai, N., Fujinuma, T., Chan, C.W., Veena and Gelvin, S.B. (2006) Constitutive expression exposes functional redundancy between the histone H2A gene and other H2A gene family members. Plant Cell 18, 1575–1589.

Yi, H.C., Mysore, K.S. and Gelvin, S.B. (2002) Expression of the Arabidopsis histone H2A-1 gene correlates with susceptibility to Agrobacterium transformation. Plant Journal 32, 285–298.

Zhou, Q., Tang, D., Huang, W., Yang, Z.M., Zhang, Y., Hamilton, J.P., Visser, R.G.F., Bachem, C.W.B., Buell, C.R., Zhang, Z.H., Zhang, C.Z. and Huang, S.W. (2020) Haplotype-resolved genome analyses of a heterozygous diploid potato. Nat Genet 52, 1018–1023.

Zhu, X.B., Chen, A.R., Butler, N.M., Zeng, Z.X., Xin, H.Y., Wang, L.X., Lv, Z.Y., Eshel, D., Douches, D.S. and Jiang, J.M. (2024) Molecular dissection of an intronic enhancer governing cold-induced expression of the vacuolar invertase gene in potato. Plant Cell 36, 1985–1999.

Zhu, Y.M., Nam, J., Humara, J.M., Mysore, K.S., Lee, L.Y., Cao, H.B., Valentine, L., Li, J.L., Kaiser, A.D., Kopecky, A.L., Hwang, H.H., Bhattacharjee, S., Rao, P.K., Tzfira, T., Rajagopal, J., Yi, H.C., Veena, Yadav, B.S., Crane, Y.M., Lin, K., Larcher, Y., Gelvin, M.J.K., Knue, M., Ramos, C., Zhao, X.W., Davis, S.J., Kim, S.I., Ranjith-Kumar, C.T., Choi, Y.J., Hallan, V.K., Chattopadhyay, S., Sui, X.Z., Ziemienowicz, A., Matthysse, A.G., Citovsky, V., Hohn, B. and Gelvin, S.B. (2003) IIdentification of Arabidopsis rat mutants. Plant Physiology 132, 494–505.

