## Supplemental Materials for "Jan and mini-Jan, a model system for potato functional genomics"

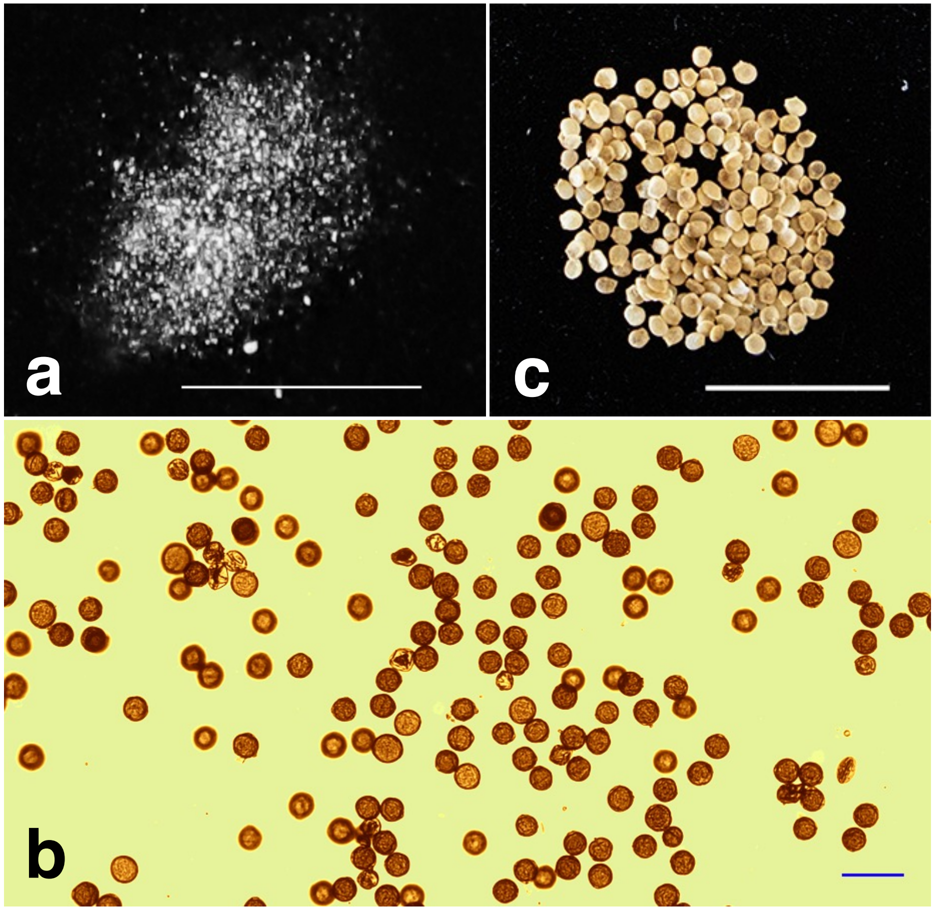


**Figure S1.** Pollen and seeds of Jan. (**a**) Pollen from a single anther. (**b**) Pollen stained by $I_2$-KI. (**c**) Seeds from two fruits. The bars represent 0.5 cm in (a) and (c), 50 µm in (b).

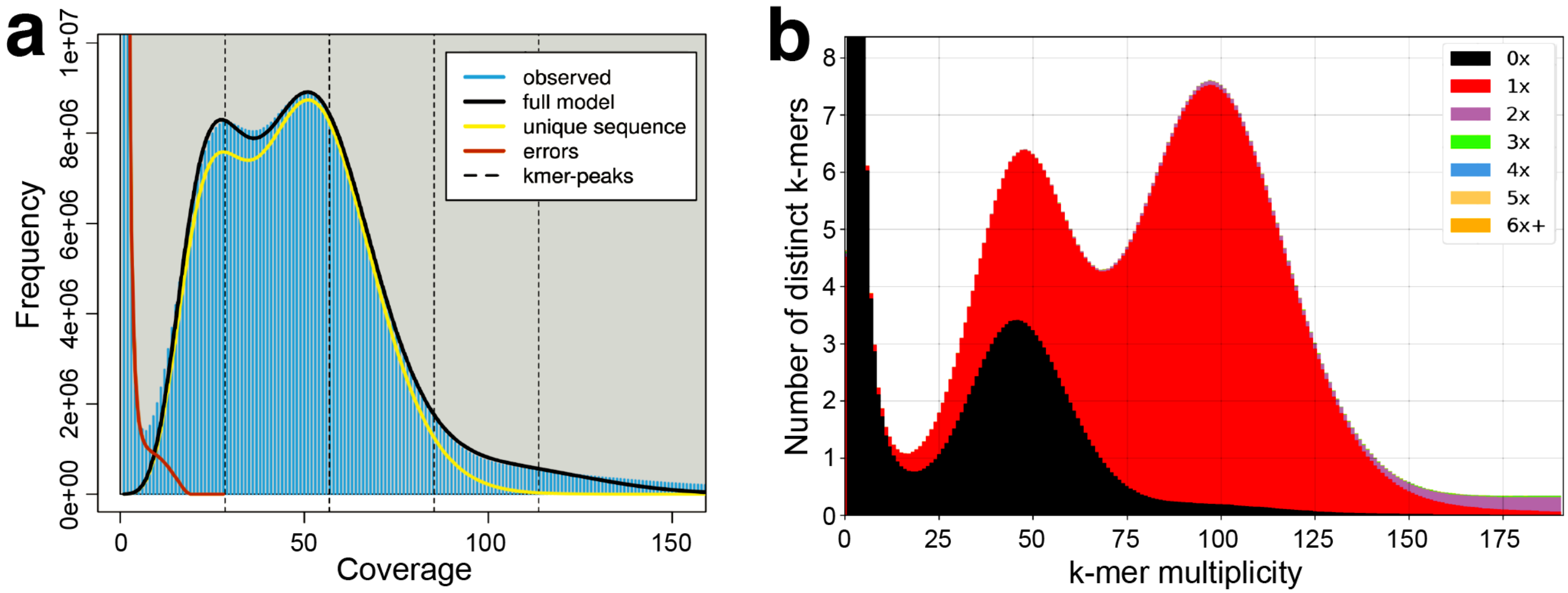


**Figure S2.** Heterozygous residue in the Jan genome. (**a**) GenomeScope plot of k-mers (k = 21) generated from Jan Illumina WGS data; the slightly bimodal distribution of k-mer frequencies indicates heterozygosity. (**b**) KAT spectra displaying presence-absence and copy numbers of WGS k-mers in Janv1.1. The black peak (k-mer multiplicity ≈ 50) represents k-mers absent in the assembly as a result of sequence deduplication, further indicative of residual heterozygosity.

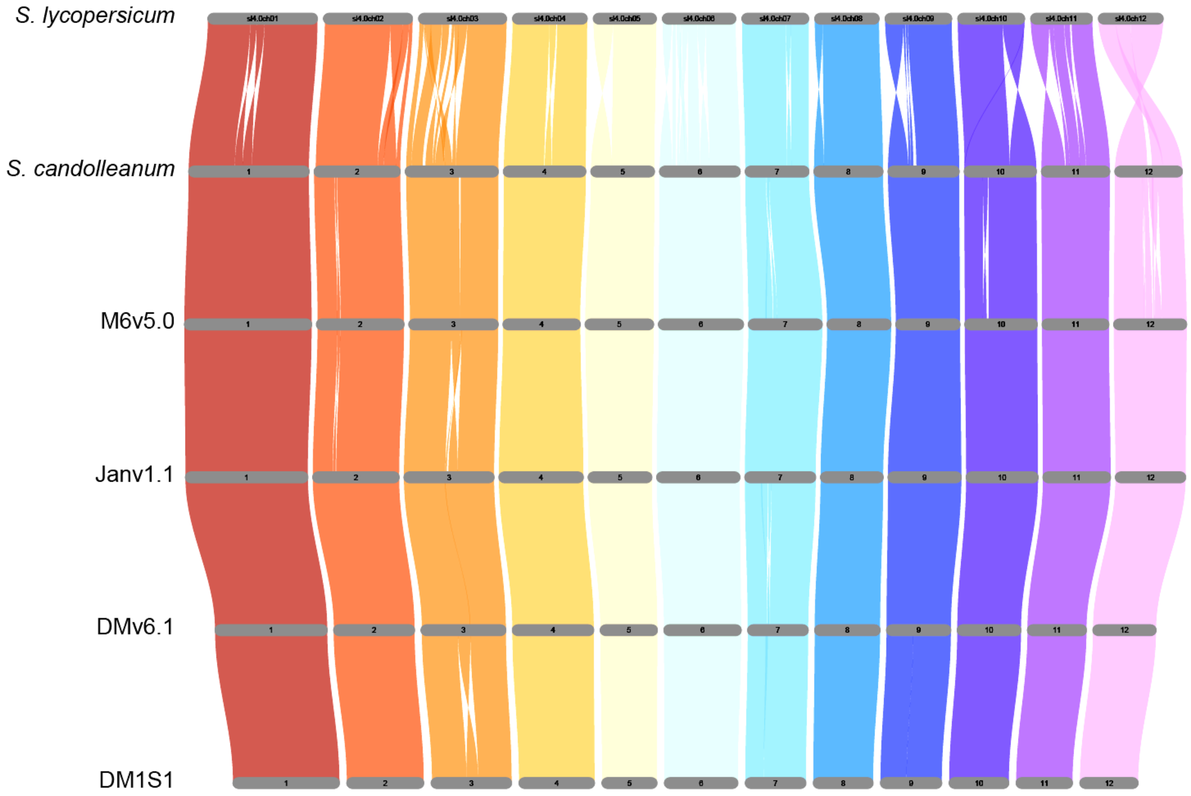


**Figure S3.** Riparian plot (scaled by number of genes) displaying genic synteny between Janv1.1, its parents (DMv6.1 and M6v5.0), other diploid potatoes (DM1S1 and *S. candolleanum*), and tomato (*S. lycopersicum*). Where available, representative high-confidence gene model sets were used as the basis for genic comparison.

#### Figure S4_a

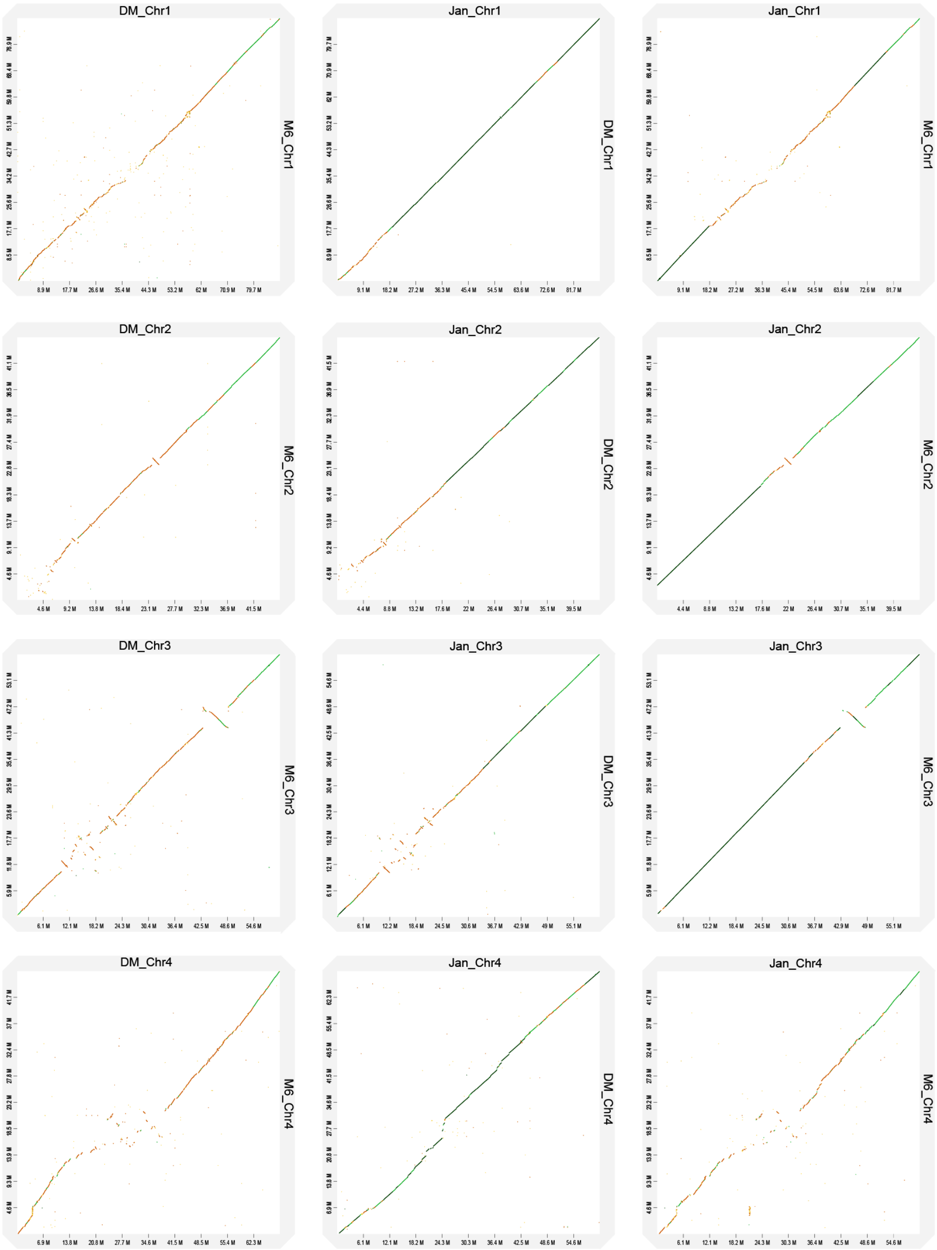
DM_Chr1
M6_Chr1
Jan_Chr1
DM_Chr1
Jan_Chr1
M6_Chr1
DM_Chr2
M6_Chr2
Jan_Chr2
DM_Chr2
Jan_Chr2
M6_Chr2
DM_Chr3
M6_Chr3
Jan_Chr3
DM_Chr3
Jan_Chr3
M6_Chr3
DM_Chr4
M6_Chr4
Jan_Chr4
DM_Chr4
Jan_Chr4
M6_Chr4

### Figure S4_b

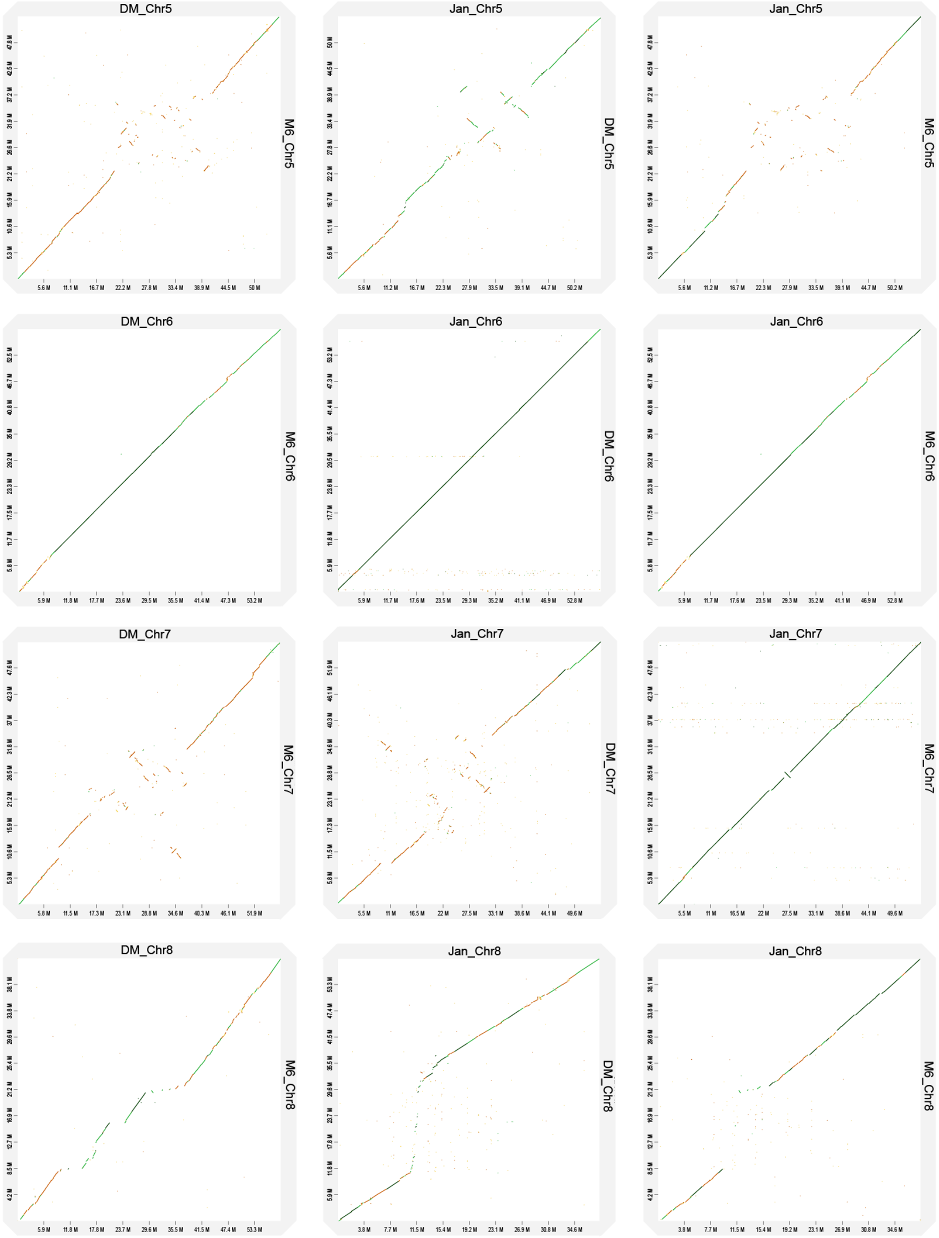

DM_Chr5
M6_Chr5
Jan_Chr5
DM_Chr5
Jan_Chr5
M6_Chr5
DM_Chr6
M6_Chr6
Jan_Chr6
DM_Chr6
Jan_Chr6
M6_Chr6
DM_Chr7
M6_Chr7
Jan_Chr7
DM_Chr7
Jan_Chr7
M6_Chr7
DM_Chr8
M6_Chr8
Jan_Chr8
DM_Chr8
Jan_Chr8
M6_Chr8

**Figure S4_c**

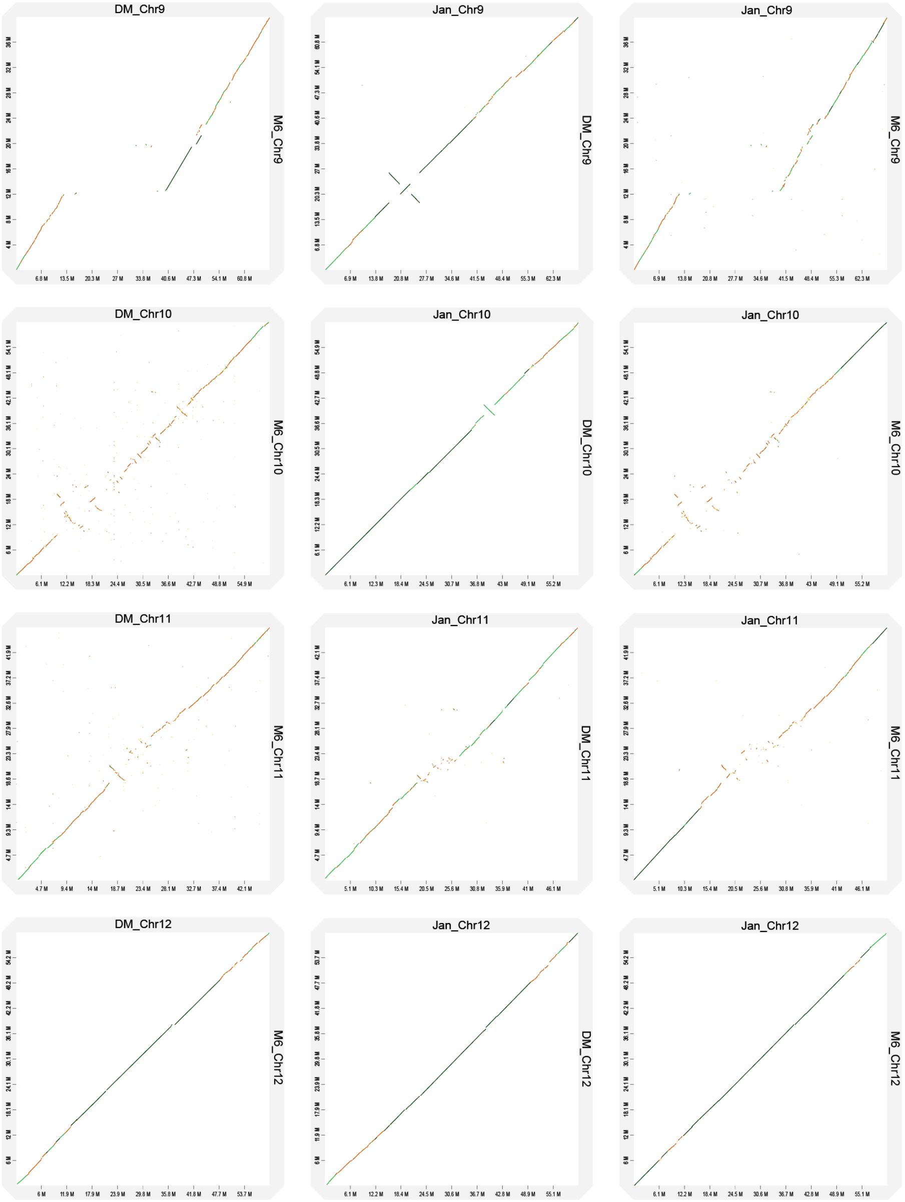


**Figure S4.** Pairwise collinearity comparisons of DMv6.1 vs. M6v5.0; Janv1.1 vs. DMv6.1; and Janv1.1 vs M6v5.0 on a chromosome-by-chromosome basis. (**a**) Comparisons of chromosomes 1-4. (**b**) Comparisons of chromosomes 5-8. (**c**) Comparisons of chromosomes 9-12.

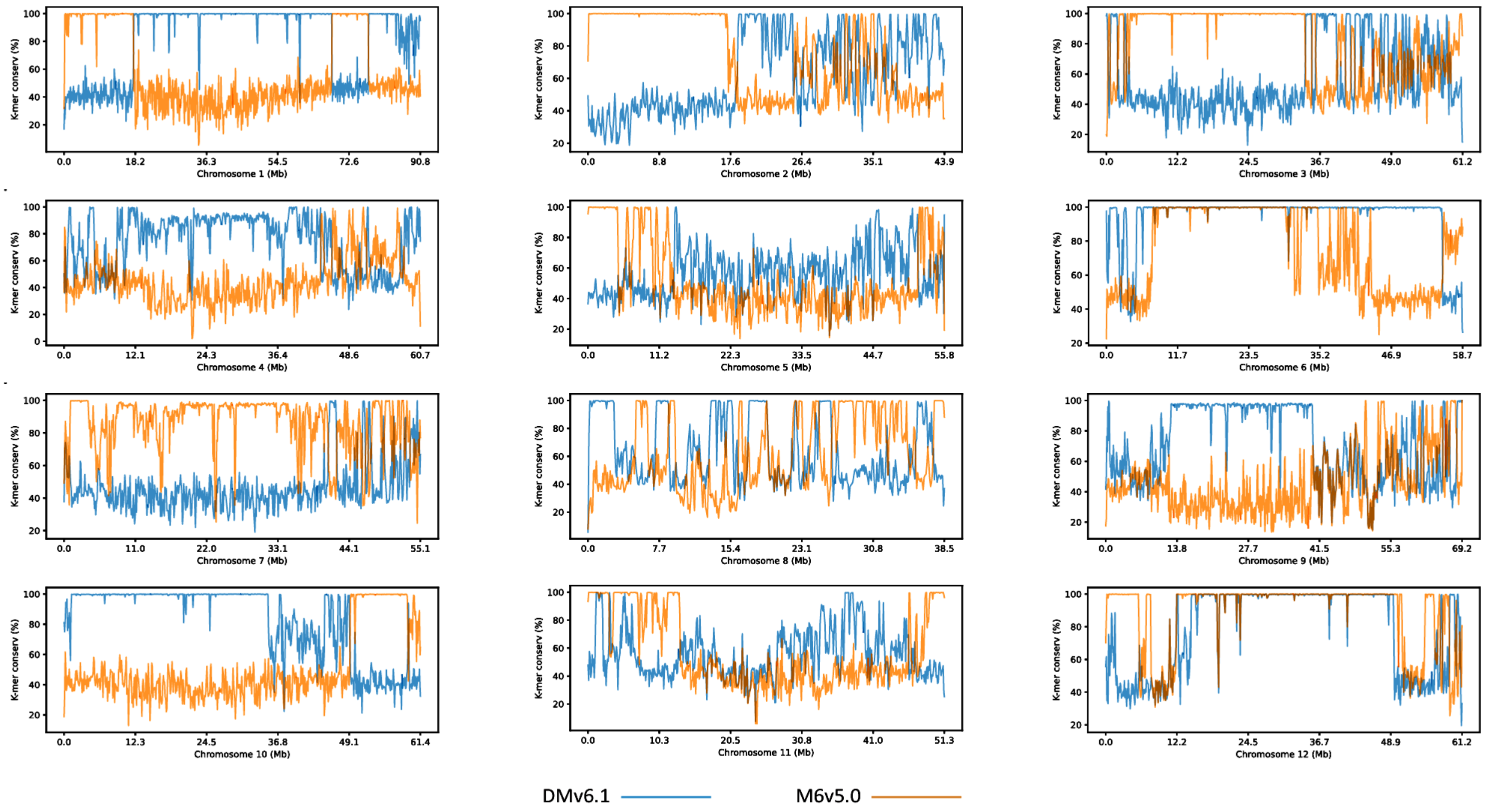


**Figure S5.** Allelic representation of DMv6.1 and M6v5.0 genomes in the Janv1.1 genome assembly. K-mers were anchored to the Janv1.1 assembly and k-mer conservation was calculated in 100 kb windows.

**Table S1.** DNA and RNA sequencing information for Jan

| SRA Run No. | Sequencing Type | Description | # Reads | Avg. read length (bp) | Total base pairs |
|---|---|---|---|---|---|
| SRR30599531 | Oxford Nanopore (ONT) | High molecular weight genomic DNA | 5,631,201 | 11,627 | 65,475,943,209 |
| SRR30810009 | Illumina | WGS DNA | 371,550,295 | 150 | 55,707,351,056 |

| SRA Run No. | Sequencing Type | Description | Tissue type | # Reads | Avg. read length (bp) | Total base pairs | Alignment rate |
|---|---|---|---|---|---|---|---|
| SRR30599841 | ONT | FL-cDNA | Immature leaf | 17,025,628 | 668 | 11,370,515,959 | - |
| SRR30599840 | ONT | FL-cDNA | Flower bud | 10,828,836 | 646 | 6,992,911,834 | - |
| SRR30599839 | ONT | FL-cDNA | Open flower | 18,100,555 | 686 | 12,411,282,203 | - |
| SRR30599838 | ONT | FL-cDNA | Tuber, flesh + peel | 11,521,741 | 774 | 8,921,064,037 | - |
| SRR30599837 | ONT | FL-cDNA | Stolon | 18,226,505 | 548 | 9,994,849,025 | - |
| SRR30599836 | ONT | FL-cDNA | Root | 22,937,208 | 684 | 15,694,158,235 | - |
| SRR30809902 | Illumina | RNA-seq | Immature leaf | 45,650,800 | 150 | 6,847,620,000 | 93.25% |
| SRR30809901 | Illumina | RNA-seq | Flower bud | 39,303,144 | 150 | 5,895,471,600 | 92.93% |
| SRR30809900 | Illumina | RNA-seq | Open flower | 46,302,471 | 150 | 6,945,370,650 | 92.63% |
| SRR30809899 | Illumina | RNA-seq | Tuber, flesh + peel | 49,000,180 | 150 | 7,350,027,000 | 90.98% |
| SRR30809898 | Illumina | RNA-seq | Stolon | 51,439,897 | 150 | 7,715,984,550 | 92.88% |
| SRR30809897 | Illumina | RNA-seq | Root | 51,825,759 | 150 | 7,773,863,850 | 92.77% |

Note: All sequence data is available under NCBI BioProject PRJNA1157315 (to be made public upon acceptance).

**Table S2.** Genome assembly summary and statistics of Jan

| Parameter | Jan: draft assembly† | Jan: contig-level assembly | ***S. tuberosum*** **Janv1.1** |
|---|---|---|---|
| Total assembly size, bp* | 920,395,782 | 717,188,417 | 717,188,417 |
| Contig N50, bp | 2,573,890 | 6,660,989 | 6,660,989 |
| Total contig No. | 2,653 | 953 | 953 |
| Scaffold N50, bp | - | - | 60,693,853 |
| Total scaffold No. | - | - | 84 |
| Chr01 length, bp | - | - | 90,828,784 |
| Chr02 length, bp | - | - | 43,902,187 |
| Chr03 length, bp | - | - | 61,239,193 |
| Chr04 length, bp | - | - | 60,832,219 |
| Chr05 length, bp | - | - | 55,912,017 |
| Chr06 length, bp | - | - | 58,696,100 |
| Chr07 length, bp | - | - | 55,166,563 |
| Chr08 length, bp | - | - | 38,538,495 |
| Chr09 length, bp | - | - | 69,240,929 |
| Chr10 length, bp | - | - | 61,364,398 |
| Chr11 length, bp | - | - | 51,297,426 |
| Chr12 length, bp | - | - | 61,243,563 |

* Total assembly size excludes Ns, which represent gaps in the assembly

† Assembly represents all contigs assembled (prior to duplication and short contig removal)

**Table S3.** BUSCO scores for Jan genome assembly and annotation

| Parameter | Assembly | Working Models | HC Models | Repr. HC Models |
|---|---|---|---|---|
| Total BUSCO groups searched | 1614 | 1614 | 1614 | 1614 |
| Complete BUSCOs (C) | 1585 (98.2%) | 1445 (89.5%) | 1441 (89.3%) | 1436 (89.0%) |
| Complete and single-copy BUSCOs (S) | 1550 (96.0%) | 828 (51.3%) | 824 (51.1%) | 1398 (86.6%) |
| Complete and duplicated BUSCOs (D) | 35 (2.2%) | 617 (38.2%) | 617 (38.2%) | 38 (2.4%) |
| Fragmented BUSCOs (F) | 5 (0.3%) | 111 (6.9%) | 109 (6.8%) | 113 (7.0%) |
| Missing BUSCOs (M) | 24 (1.5%) | 58 (3.6%) | 64 (3.9%) | 65 (4.0%) |

Note: Assessment of Jan genome assembly and annotation completeness with Benchmarking Universal Single-Copy Orthologs (BUSCO) analysis (v5.4.3; Simão et al. 2015; Manni et al. 2021) with embryophyta_odb10 set as the lineage dataset. For completeness analysis of the assembly, BUSCO was run in eukaryotic genome mode using the gene predictor Metaeuk (v7.bba0d80, https://github.com/soedinglab/metaeuk) (Levy Karin et al. 2020). For completeness analysis of the annotation (performed for all working gene models, high-confidence (HC) gene models, and representative HC gene models), BUSCO was run in "proteins" mode.

**Table S4.** Repetitive DNA in the Jan genome

| | | | |
|---|---|---|---|
| Sequences: | 84 | | |
| Total length: | 717275317 bp (717188417 bp excl N/X-runs) | | |
| GC level: | 34.81% | | |
| Bases masked: | 471892781 bp (65.79%) | | |
| **Element type** | **Number of elements*** | **Length occupied (bp)** | **Percentage of sequence** |
| Retroelements | 187622 | 189905019 | 26.48% |
| SINEs | 2304 | 282885 | 0.04% |
| Penelope | 249 | 69379 | 0.01% |
| LINEs | 39665 | 16567018 | 2.31% |
| CRE/SLACS | 0 | 0 | 0.00% |
| L2/CR1/Rex | 0 | 0 | 0.00% |
| R1/LOA/Jockey | 0 | 0 | 0.00% |
| R2/R4/NeSL | 0 | 0 | 0.00% |
| RTE/Bov-B | 17291 | 4324236 | 0.60% |
| L1/CIN4 | 22203 | 12209695 | 1.70% |
| LTR elements | 145653 | 173055116 | 24.13% |
| BEL/Pao | 217 | 22134 | 0.00% |
| Ty1/Copia | 33195 | 24089868 | 3.36% |
| Gypsy/DIRS1 | 105571 | 138283821 | 19.28% |
| Retroviral | 654 | 324013 | 0.05% |
| DNA transposons | 26331 | 14559365 | 2.03% |
| hobo-Activator | 5888 | 3704237 | 0.52% |
| Tc1-IS630-Pogo | 1150 | 317451 | 0.04% |
| En-Spm | 0 | 0 | 0.00% |
| MULE-MuDR | 10426 | 5416522 | 0.76% |
| PiggyBac | 0 | 0 | 0.00% |
| Tourist/Harbinger | 3283 | 1056237 | 0.15% |
| Other (Mirage, P-element, Transib) | 0 | 0 | 0.00% |
| Rolling-circles | 4866 | 2464750 | 0.34% |
| Unclassified | 833071 | 256709566 | 35.79% |
| Total interspersed repeats: | | 461243329 | 64.30% |
| Small RNA | 1844 | 228167 | 0.03% |
| Satellites | 271 | 145750 | 0.02% |
| Simple repeats | 108390 | 5830500 | 0.81% |
| Low complexity | 22184 | 2004661 | 0.28% |

* Most repeats fragmented by insertions or deletions have been counted as one element.

**Table S5.** GO terms enriched in Jan genes inherited from DM and M6

**Genes inherited from DM (Biological Process)**

| GO Term | Name | # Significant Genes | *p*-value |
|---|---|---|---|
| GO:0002376 | immune system process | 10 | 0.00523 |
| GO:0006022 | aminoglycan metabolic process | 10 | 0.00065 |
| GO:0006026 | aminoglycan catabolic process | 10 | 0.00065 |
| GO:0006040 | amino sugar metabolic process | 10 | 0.00119 |
| GO:0006091 | generation of precursor metabolites and energy | 93 | 0.0042 |
| GO:0006412 | translation | 206 | 0.00836 |
| GO:0006508 | proteolysis | 297 | 0.00069 |
| GO:0006576 | biogenic amine metabolic process | 20 | 0.00415 |
| GO:0006629 | lipid metabolic process | 318 | 0.0034 |
| GO:0006721 | terpenoid metabolic process | 37 | 0.00415 |
| GO:0006779 | porphyrin-containing compound biosynthetic process | 25 | 0.00013 |
| GO:0006955 | immune response | 10 | 0.00523 |
| GO:0009056 | catabolic process | 266 | 8.30E-06 |
| GO:0009057 | macromolecule catabolic process | 135 | 0.00026 |
| GO:0009605 | response to external stimulus | 65 | 0.00562 |
| GO:0009607 | response to biotic stimulus | 40 | 0.00863 |
| GO:0009620 | response to fungus | 20 | 0.00827 |
| GO:0009832 | plant-type cell wall biogenesis | 13 | 0.00131 |
| GO:0010154 | fruit development | 9 | 0.00537 |
| GO:0010383 | cell wall polysaccharide metabolic process | 27 | 0.00124 |
| GO:0015979 | photosynthesis | 39 | 0.00266 |
| GO:0016050 | vesicle organization | 11 | 0.00022 |
| GO:0019684 | photosynthesis, light reaction | 25 | 0.00033 |
| GO:0022900 | electron transport chain | 25 | 0.00866 |
| GO:0030026 | intracellular manganese ion homeostasis | 9 | 0.00189 |
| GO:0032787 | monocarboxylic acid metabolic process | 84 | 0.00668 |
| GO:0032879 | regulation of localization | 11 | 0.00022 |
| GO:0043043 | peptide biosynthetic process | 211 | 0.00462 |
| GO:0044036 | cell wall macromolecule metabolic process | 37 | 1.10E-05 |
| GO:0044248 | cellular catabolic process | 124 | 0.00239 |
| GO:0044419 | biological process involved in interspecies interaction between organisms | 40 | 0.00983 |
| GO:0045040 | protein insertion into mitochondrial outer membrane | 7 | 0.00241 |
| GO:0050832 | defense response to fungus | 20 | 0.00827 |
| GO:0051049 | regulation of transport | 11 | 0.00022 |
| GO:0051649 | establishment of localization in cell | 187 | 0.00941 |
| GO:0055071 | manganese ion homeostasis | 9 | 0.00189 |

| | | | |
|---|---|---|---|
| GO:0071554 | cell wall organization or biogenesis | 95 | 1.20E-07 |
| GO:1901071 | glucosamine-containing compound metabolic process | 10 | 0.00065 |
| GO:1901565 | organonitrogen compound catabolic process | 119 | 0.00108 |

**Genes inherited from DM (Molecular Function)**

| GO Term | Name | # Significant genes | *p*-value |
|---|---|---|---|
| GO:0000287 | magnesium ion binding | 86 | 0.00552 |
| GO:0000976 | transcription cis-regulatory region binding | 38 | 0.00632 |
| GO:0001067 | transcription regulatory region nucleic acid binding | 38 | 0.00632 |
| GO:0001216 | DNA-binding transcription activator activity | 11 | 0.00597 |
| GO:0001228 | DNA-binding transcription activator activity, RNA polymerase II-specific | 11 | 0.00597 |
| GO:0003677 | DNA binding | 677 | 0.00419 |
| GO:0003735 | structural constituent of ribosome | 160 | 9.80E-05 |
| GO:0004325 | ferrochelatase activity | 9 | 0.00121 |
| GO:0004497 | monooxygenase activity | 192 | 1.10E-06 |
| GO:0004523 | RNA-DNA hybrid ribonuclease activity | 49 | 0.00206 |
| GO:0004568 | chitinase activity | 10 | 0.00079 |
| GO:0004857 | enzyme inhibitor activity | 102 | 7.50E-06 |
| GO:0005198 | structural molecule activity | 217 | 6.20E-05 |
| GO:0005506 | iron ion binding | 209 | 2.60E-05 |
| GO:0008061 | chitin binding | 21 | 5.20E-05 |
| GO:0008094 | ATP-dependent activity, acting on DNA | 75 | 0.00284 |
| GO:0008171 | O-methyltransferase activity | 45 | 0.00015 |
| GO:0008234 | cysteine-type peptidase activity | 68 | 0.00113 |
| GO:0008417 | fucosyltransferase activity | 6 | 0.00853 |
| GO:0008970 | phospholipase A1 activity | 20 | 1.00E-07 |
| GO:0009044 | xylan 1,4-beta-xylosidase activity | 9 | 0.00225 |
| GO:0009055 | electron transfer activity | 45 | 0.00058 |
| GO:0010333 | terpene synthase activity | 52 | 1.70E-05 |
| GO:0016298 | lipase activity | 38 | 0.00288 |
| GO:0016491 | oxidoreductase activity | 702 | 0.00102 |
| GO:0016705 | oxidoreductase activity, acting on paired donors, with incorporation or reduction of molecular oxygen | 209 | 2.10E-08 |
| GO:0016717 | oxidoreductase activity, acting on paired donors, with oxidation of a pair of donors resulting in the reduction of molecular oxygen to two molecules of water | 12 | 0.00256 |
| GO:0016835 | carbon-oxygen lyase activity | 94 | 0.00022 |
| GO:0016838 | carbon-oxygen lyase activity, acting on phosphates | 55 | 6.20E-05 |
| GO:0019199 | transmembrane receptor protein kinase activity | 10 | 0.00402 |
| GO:0020037 | heme binding | 244 | 2.10E-07 |
| GO:0022804 | active transmembrane transporter activity | 199 | 0.00382 |

| | | | |
|---|---|---|---|
| GO:0030145 | manganese ion binding | 50 | 8.90E-12 |
| GO:0030527 | structural constituent of chromatin | 29 | 0.00053 |
| GO:0042626 | ATPase-coupled transmembrane transporter activity | 100 | 0.00242 |
| GO:0045735 | nutrient reservoir activity | 9 | 0.00632 |
| GO:0046906 | tetrapyrrole binding | 249 | 6.50E-08 |
| GO:0046915 | transition metal ion transmembrane transporter activity | 16 | 0.00698 |
| GO:0046982 | protein heterodimerization activity | 53 | 0.00612 |

**Genes inherited from M6 (Biological Process)**

| GO Term | Name | # Significant genes | *p*-value |
|---|---|---|---|
| GO:0000724 | double-strand break repair via homologous recombination | 9 | 0.00434 |
| GO:0006412 | translation | 219 | 4.20E-05 |
| GO:0006476 | protein deacetylation | 20 | 0.0017 |
| GO:0006518 | peptide metabolic process | 226 | 0.00034 |
| GO:0006631 | fatty acid metabolic process | 48 | 0.00648 |
| GO:0006869 | lipid transport | 49 | 0.00131 |
| GO:0006950 | response to stress | 404 | 5.00E-05 |
| GO:0006952 | defense response | 186 | 0.00863 |
| GO:0006979 | response to oxidative stress | 93 | 4.60E-08 |
| GO:0008037 | cell recognition | 47 | 1.30E-08 |
| GO:0009611 | response to wounding | 21 | 3.00E-06 |
| GO:0009627 | systemic acquired resistance | 14 | 0.00038 |
| GO:0009664 | plant-type cell wall organization | 27 | 0.00189 |
| GO:0009719 | response to endogenous stimulus | 123 | 0.00056 |
| GO:0009725 | response to hormone | 122 | 0.00068 |
| GO:0010876 | lipid localization | 51 | 0.00072 |
| GO:0015711 | organic anion transport | 29 | 0.00305 |
| GO:0015849 | organic acid transport | 27 | 0.0006 |
| GO:0015986 | proton motive force-driven ATP synthesis | 17 | 0.00194 |
| GO:0022618 | protein-RNA complex assembly | 22 | 0.00955 |
| GO:0022622 | root system development | 22 | 7.30E-05 |
| GO:0032502 | developmental process | 111 | 0.00531 |
| GO:0032787 | monocarboxylic acid metabolic process | 82 | 0.00635 |
| GO:0033993 | response to lipid | 40 | 0.00586 |
| GO:0035601 | protein deacylation | 20 | 0.0017 |
| GO:0042221 | response to chemical | 178 | 0.00444 |
| GO:0042273 | ribosomal large subunit biogenesis | 9 | 0.00434 |
| GO:0042744 | hydrogen peroxide catabolic process | 73 | 7.40E-13 |
| GO:0043043 | peptide biosynthetic process | 219 | 9.50E-05 |
| GO:0043603 | amide metabolic process | 249 | 0.00022 |

| | | | |
|---|---|---|---|
| GO:0043604 | amide biosynthetic process | 233 | 6.00E-05 |
| GO:0044248 | cellular catabolic process | 128 | 0.00016 |
| GO:0045229 | external encapsulating structure organization | 54 | 0.00028 |
| GO:0046942 | carboxylic acid transport | 27 | 0.0006 |
| GO:0048364 | root development | 22 | 7.30E-05 |
| GO:0048544 | recognition of pollen | 47 | 1.30E-08 |
| GO:0048856 | anatomical structure development | 94 | 0.00594 |
| GO:0050896 | response to stimulus | 679 | 0.00167 |
| GO:0051259 | protein complex oligomerization | 10 | 0.00265 |

**Genes inherited from M6 (Molecular Function)**

| GO Term | Name | # Significant genes | *p*-value |
|---|---|---|---|
| GO:0003735 | structural constituent of ribosome | 165 | 1.00E-07 |
| GO:0003824 | catalytic activity | 3539 | 0.00241 |
| GO:0004072 | aspartate kinase activity | 10 | 0.00077 |
| GO:0004175 | endopeptidase activity | 135 | 0.00124 |
| GO:0004412 | homoserine dehydrogenase activity | 7 | 0.00345 |
| GO:0004497 | monooxygenase activity | 207 | 4.70E-13 |
| GO:0004527 | exonuclease activity | 32 | 4.90E-05 |
| GO:0004857 | enzyme inhibitor activity | 106 | 1.20E-08 |
| GO:0005198 | structural molecule activity | 232 | 2.20E-10 |
| GO:0005199 | structural constituent of cell wall | 11 | 0.00133 |
| GO:0005388 | P-type calcium transporter activity | 13 | 0.00257 |
| GO:0005506 | iron ion binding | 214 | 1.00E-08 |
| GO:0008194 | UDP-glycosyltransferase activity | 183 | 7.00E-10 |
| GO:0016209 | antioxidant activity | 104 | 3.50E-08 |
| GO:0016491 | oxidoreductase activity | 719 | 3.90E-10 |
| GO:0016679 | oxidoreductase activity, acting on diphenols and related substances as donors | 22 | 5.00E-05 |
| GO:0016682 | oxidoreductase activity, acting on diphenols and related substances as donors, oxygen as acceptor | 22 | 3.40E-05 |
| GO:0016684 | oxidoreductase activity, acting on peroxide as acceptor | 95 | 5.40E-10 |
| GO:0016705 | oxidoreductase activity, acting on paired donors, with incorporation or reduction of molecular oxygen | 203 | 7.20E-10 |
| GO:0016757 | glycosyltransferase activity | 307 | 7.20E-12 |
| GO:0016774 | phosphotransferase activity, carboxyl group as acceptor | 11 | 0.00938 |
| GO:0016841 | ammonia-lyase activity | 11 | 0.00471 |
| GO:0016896 | RNA exonuclease activity, producing 5-prime-phosphomonoesters | 13 | 0.00257 |
| GO:0017171 | serine hydrolase activity | 93 | 0.00884 |
| GO:0019208 | phosphatase regulator activity | 19 | 0.0042 |
| GO:0019840 | isoprenoid binding | 9 | 0.00032 |

| | | | |
|---|---|---|---|
| GO:0019888 | protein phosphatase regulator activity | 18 | 0.00793 |
| GO:0020037 | heme binding | 277 | 3.80E-19 |
| GO:0030246 | carbohydrate binding | 100 | 4.00E-05 |
| GO:0030247 | polysaccharide binding | 53 | 1.40E-07 |
| GO:0030527 | structural constituent of chromatin | 33 | 1.90E-06 |
| GO:0033293 | monocarboxylic acid binding | 12 | 0.00016 |
| GO:0035251 | UDP-glucosyltransferase activity | 31 | 0.00954 |
| GO:0042562 | hormone binding | 10 | 0.00077 |
| GO:0042887 | amide transmembrane transporter activity | 21 | 0.00864 |
| GO:0043178 | alcohol binding | 9 | 0.00222 |
| GO:0046906 | tetrapyrrole binding | 279 | 4.60E-19 |
| GO:0061134 | peptidase regulator activity | 39 | 0.00396 |
| GO:0098772 | molecular function regulator activity | 185 | 0.00042 |
